# Disrupting the Transmembrane Domain Oligomerization of Protein Tyrosine Phosphatase Receptor J Promotes its Activity in Cancer Cells

**DOI:** 10.1101/672147

**Authors:** Elizabeth Bloch, Eden L. Sikorski, David Pontoriero, Evan K. Day, Bryan W. Berger, Matthew J. Lazzara, Damien Thévenin

## Abstract

Despite the critical regulatory roles that receptor protein tyrosine phosphatases (RPTP) play in mammalian signal transduction, the detailed structural basis for the regulation of their catalytic activity is not fully understood, nor are they generally therapeutically targetable. It is due, in part, to the lack of known natural ligands or selective agonists. In contrast to conventional structure-function relationship for receptor tyrosine kinases (RTKs), the activity of RPTPs has been reported to be suppressed by dimerization, which may prevent their access to their RTK substrates. We report here that: (i) homodimerization of PTPRJ (also known as DEP-1) is regulated by specific transmembrane (TM) residues, and (ii) disrupting these interactions can destabilize full-length PTPRJ homodimerization in cells, reduce the phosphorylation of EGFR (a known substrate) and downstream signaling effectors, antagonize EGFR-driven cell phenotypes, and promote substrate access. We demonstrate these points in human cancer cells using both mutational studies and through the identification of a peptide designed to bind to the PTPRJ TM domain. This peptide is the first example of such allosteric agonist of RPTPs. This study, therefore, provides not only fundamental structure-function insights on how PTPRJ activity is tuned by TM interactions in cells but also opportunities to develop a unique class of agents that could be used as tools to probe RPTPs signaling regulating mechanisms or for therapeutic purposes in cancers driven by RTK signaling.

## Introduction

Normally, cell signaling by receptor tyrosine kinases (RTKs) is tightly controlled by the counterbalancing actions of protein tyrosine phosphatases (PTPs), which dephosphorylate regulatory tyrosines on RTKs to attenuate their signal-initiating potency. Twenty-two PTPs are classified as receptor-like PTPs (RPTPs), which all share the same architecture: an extracellular region (often including fibronectin type III repeats and immunoglobin-like domains), a single-pass transmembrane segment, and either one or two cytosolic highly conserved PTP domains (1, 2). These integral membrane PTPs are some of the most important regulators of RTKs, and their normal function is critical for homeostatic control (2, 3). Interestingly, while RPTPs can act as suppressors of several tumors, including colon, lung, breast, and thyroid cancers, they can also act as oncogenes when deregulated either through reduced expression, often due to loss of heterozygosity or loss-of-function mutations (4-8). Notably, fewer than one-third of reported point mutations occur within the intracellular phosphatase domains indicating inactivation occurs without structural changes to the catalytic domain (9). RPTPs have therefore long been viewed as potential therapeutic targets.

Of the RPTPs characterized, PTPRJ (also known as DEP-1) is particularly intriguing as it has been identified as a regulator of C-terminal tyrosine phosphorylation in the epidermal growth factor receptor (EGFR) (10-12), a frequent driver of oncogenic signaling across numerous cancers. Deregulation of mechanisms controlling EGFR activation and expression results in abnormal cell growth and proliferation. PTPRJ has a direct role in regulating EGFR activity, implicating it as a relevant tumor suppressor. Furthermore, PTPRJ is often down-regulated in numerous cancers, and its restoration was shown to inhibit malignant phenotypes both *in vitro* and *in vivo* (13).

While small molecule allosteric inhibition of some non-receptor PTPs is now possible (14), methods to specifically target RPTPs are missing, due, in part, to the limited understanding of their mechanism of action and the lack of known natural ligands. Indeed, compared to the structure-function relationships for RTKs, relatively little has been elucidated for RPTPs. For instance, the reported ability of homodimerization to antagonize RPTP catalytic activity appears to be a general feature of the entire family, but there is no proposed common model. The “head-to-toe” dimerization model for the PTP D1 and D2 intracellular domains, proposed originally by Barr *et al.* for PTPRG, is generally accepted for RPTPs with tandemly repeated intracellular domains (1). However, since PTPRJ and other members of the R3 sub-family have only one catalytic intracellular PTP domain, the “head-to-toe” model and the “inhibitory wedge” model proposed for PTPRA (15, 16) seem to be incompatible. Nevertheless, the reported ability of homodimerization to antagonize PTPRJ catalytic activity and substrate access presents potential opportunities to develop strategies to promote RPTP activity against their oncogenic RTK substrate (17). While the transmembrane (TM) domain of several RPTPs has been proposed to be involved in their homodimerization (17-20), and therefore represents an attractive target, there is no clear structure-based proposal for how this occurs. Therefore, elucidating the contribution of the TM domain in RPTPs, and particularly in PTPRJ regulation, can provide significant insight into how these receptors function, interact, and eventually be modulated, leading to new methods to target signaling of oncogenic RTKs that may be less susceptible to common mechanisms of resistance.

Here, we employed mutational studies to show that PTPRJ homodimerization is regulated through specific TM residue interactions. Furthermore, disrupting these interactions antagonizes PTPRJ homodimerization, thereby promoting its phosphatase activity and ultimately inhibiting EGFR-driven cell phenotypes. We used these new findings to identify and characterize a synthetic peptide that interacts with and disrupts PTPRJ homodimers through specific TM interactions. We show the delivery of this peptide selectively modulates the dimerization state and activity of PTPRJ in EGFR-driven cancer cells. The present study represents a structure-function determination of the TM domain of PTPRJ, and a new way to selectively modulate the activity of this important class of phosphatases in cancer cells.

## Results

### PTPRJ self-association is mediated by specific TM residues

To assess whether the transmembrane (TM) domain and specific amino acid residues play a role in the self-association of PTPRJ, we first used the dominant-negative AraC-based transcriptional reporter assay (DN-AraTM) (21, 22). This assay reports on the propensity of TM domains to self-associate and heterodimerize in cell membranes. Briefly, it relies on a protein chimera containing the receptor domain of interest fused to either the transcription factor AraC (which is active at the ara*BAD* promoter as a homodimer), or to an AraC mutant unable to activate transcription (AraC*). Both chimeras include an N-terminal maltose-binding protein (MBP) fusion that directs chimera insertion in the inner membrane of AraC-deficient *E. coli* (SB1676). Homodimerization of AraC (a result from receptor domain self-association) induces the transcription of the gene coding for the green fluorescent protein (GFP). Thus, GFP fluorescence intensity is a measure of receptor domain dimerization.

When the TM domain of wild-type PTPRJ (WT) (Figure 1A) was expressed as a fusion to AraC, a strong GFP signal was observed, indicating that the TM domain has a high propensity to self-associate (Figure 1B; black bar). On the other hand, when PTPRJ WT-AraC* was co-expressed as a competitor to PTPRJ WT-AraC, a significant decrease in GFP signal was observed (Figure 1B, hashed bar), consistent with specific TM-TM interactions. Having confirmed that PTPRJ self-associates, in part through the TM domain, we sought to identify the amino acids required for this interaction. To alter interfacial packing and introduce steric clashes, amino acids in the TM region with small side chains (Ala, Cys, Gly,) were mutated to leucine, while those with larger side chains (Ile, Leu, Phe, Val) were mutated to alanine (23). The propensity of each construct to self-associate was then analyzed using the DN-AraTM assay. Among all the tested TM residues, three mutations (G979L, G983L, and G983L) caused the greatest decrease in GFP signal compared to WT (Figure 1C), suggesting that these three residues are involved in stabilizing PTPRJ self-association. Interestingly, the arrangement of these three glycines places them on the same helix face (Figure 1D) and is consistent with a double GxxxG zipper (18), a common motif for TM packing interfaces (24-28), and further pointing to specific homodimeric interactions. Immunoblotting and maltose complementation assays showed that all constructs were expressed at similar levels (Figure 1C, inset) and properly inserted in the membrane (Figure S1), confirming that variation in the fluorescence signal was solely due to changes in association state of the constructs.

**Figure 1.**
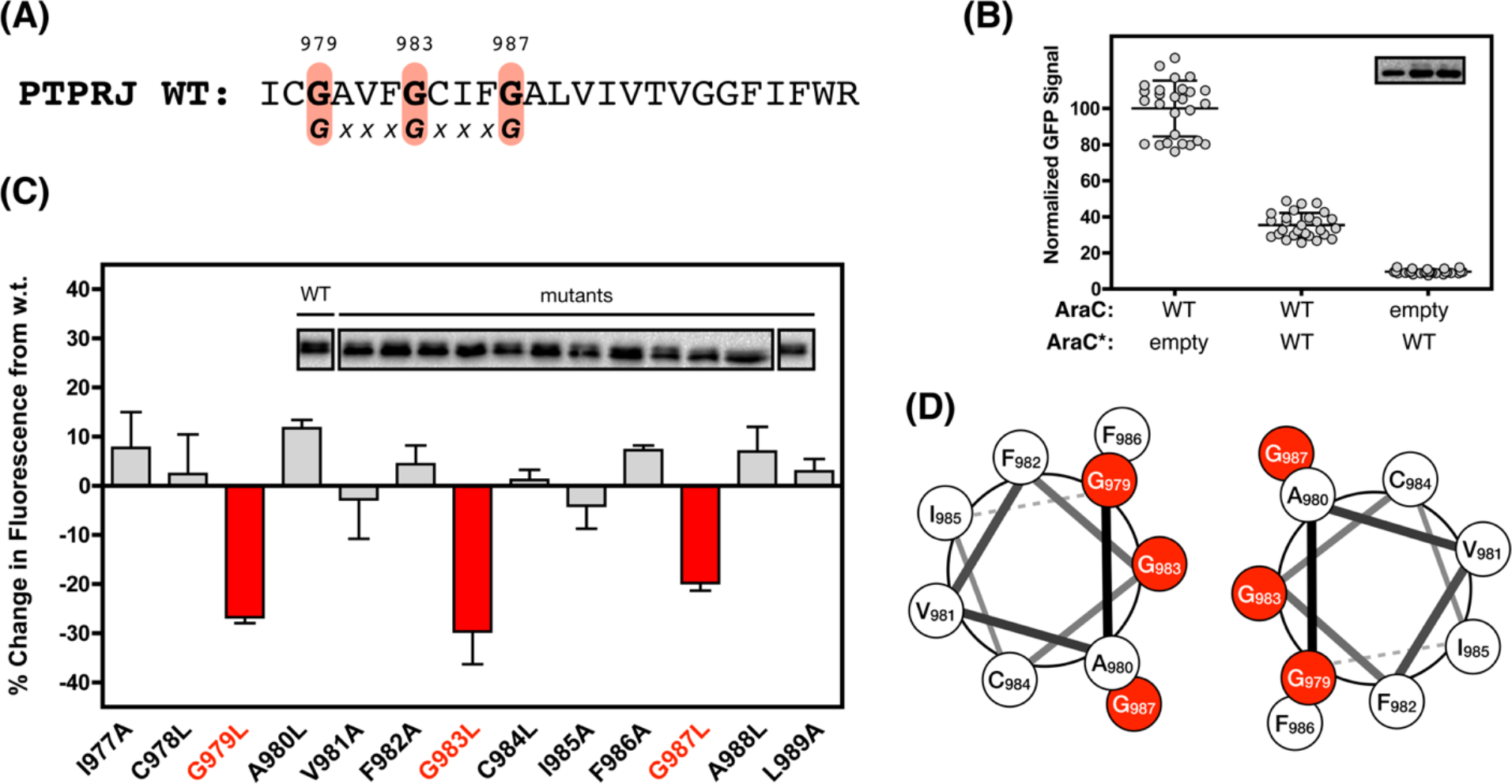
PTPRJ self-associates through specific TM residues. **(A)** Sequence of the TM domain of wild-type PTPRJ (WT). (red: glycine residues part of the double GxxxG motif). **(B)** GFP signal measured using the DN-AraTM assay with wild-type PTPRJ. Results are shown as mean ± SEM (*n* = 9). **(B, inset)** Representative immunoblot against MBP (from the same gel). **(C)** AraTM results of Ala-Leu mutagenesis scan normalized to WT PTPRJ. Results are shown as mean ± SEM (*n* = 9). **(C, inset)** Representative immunoblot against MBP, showing similar expression between WT and mutants. **(D)** Helical wheel diagram representation of the possible PTPRJ helix dimer mediated by the double GxxxG motif (red)

### PTPRJ point mutants decrease EGFR phosphorylation, downstream signaling, and inhibit EGFR-driven cell migration

Next, we determined the functional significance of receptor point mutations in cells. We hypothesized that disrupting PTPRJ self-association through point mutations at G979, G983, and G983 would lead to a decrease in phosphorylation of EGFR and downstream signaling intermediates, and ultimately inhibit EGFR-driven cancer cell migration. To test our hypothesis, full-length WT, G979L, G983L and G987L PTPRJ were ectopically and stably expressed in UMSCC2 cells, an EGFR-driven human squamous cell carcinoma cell line that exhibits very little PTPRJ expression (Figure 2A; EV). All four cell lines expressed similar levels of PTPRJ and EGFR (Figure S2). After serum starvation and treatment with EGF, the level of phosphorylation of EGFR (at Y1068 and Y1173, two representatives of EGFR phosphorylation across cytoplasmic tyrosines) (*28*, *29*), and of ERK1/2 was measured by immunoblotting. Consistent with our hypothesis, a significant decrease in EGFR and ERK1/2 phosphorylation was observed with the glycine to leucine point mutants when compared to wild-type PTPRJ (Figure 2; + 10 ng/mL EGF). Moreover, not only does expression of WT PTPRJ significantly inhibit EGFR-driven cell migration when compared to empty vector (Figure 3), but G983L and G987L receptor mutants also inhibit cell migration to an even greater extent than WT PTPRJ (Figure 3). Remarkably, these results are consistent with the immunoblotting data, where G983L and G987L receptor mutants lowered EGFR phosphorylation the most (Figure 2).

**Figure 2.**
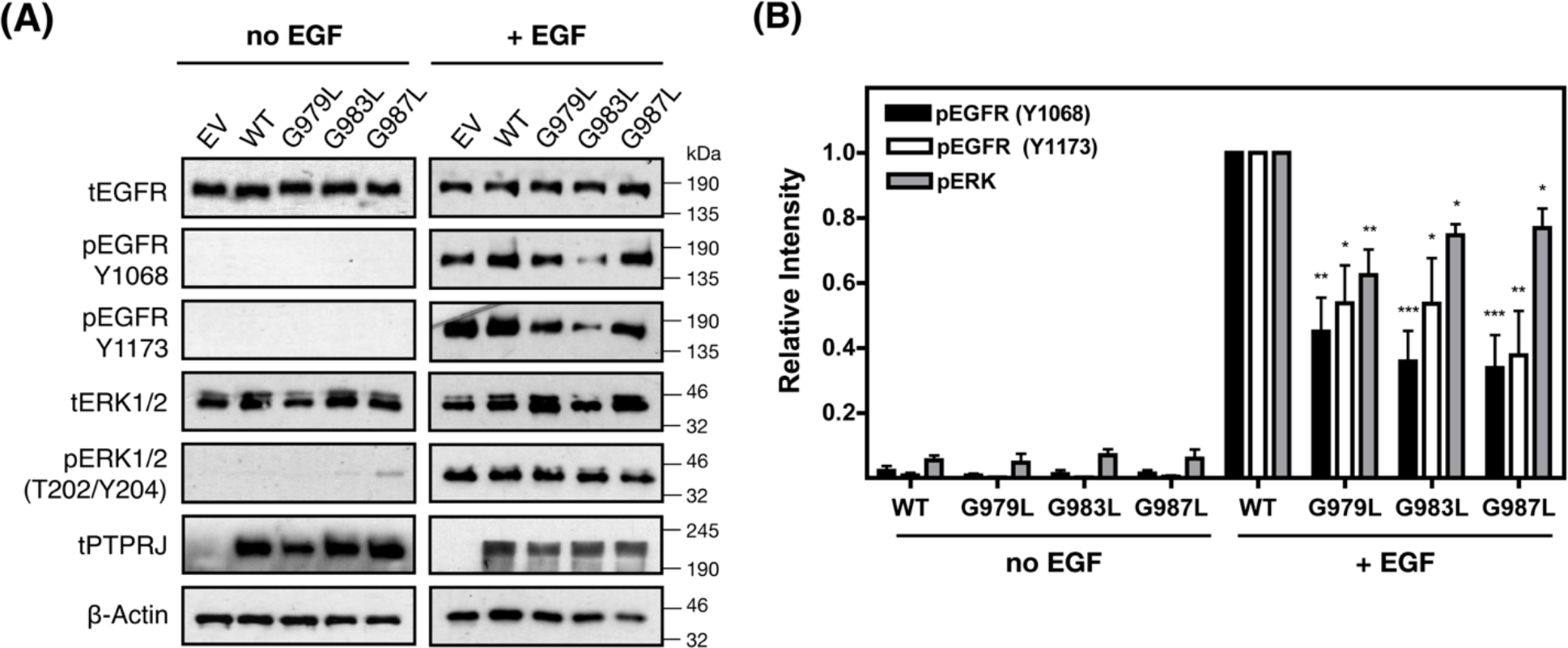
PTPRJ point mutants decrease phosphorylation of EGFR and ERK. Representative immunoblot **(A)** and quantification **(B)** of the effect of PTPRJ transmembrane mutations on EGFR and ERK1/2 phosphorylation levels. Stimulated (+ EGF) and unstimulated (-EGF) serum-starved UMSCC2 cells were probed for EGFR, phospho-EGFR (Y1068 and Y1173), ERK1/2, phospho-ERK1/2 (T202/Y204) and PTPRJ by immunoblot. Representative blots from the same samples are shown. **(B)** The relative (ratio of phosphorylated to total protein) intensities are shown as mean ± SEM (*n* = 3-5). Statistical significance between WT and mutants (for each phosphorylation site) was assessed using one-way ANOVA corrected for multiple comparison (Dunnet test) at 95% confidence intervals: \*\*\**p* ≤0.001; \*\**p* ≤ 0.01; and \**p* ≤ 0.05

**Figure 3.**
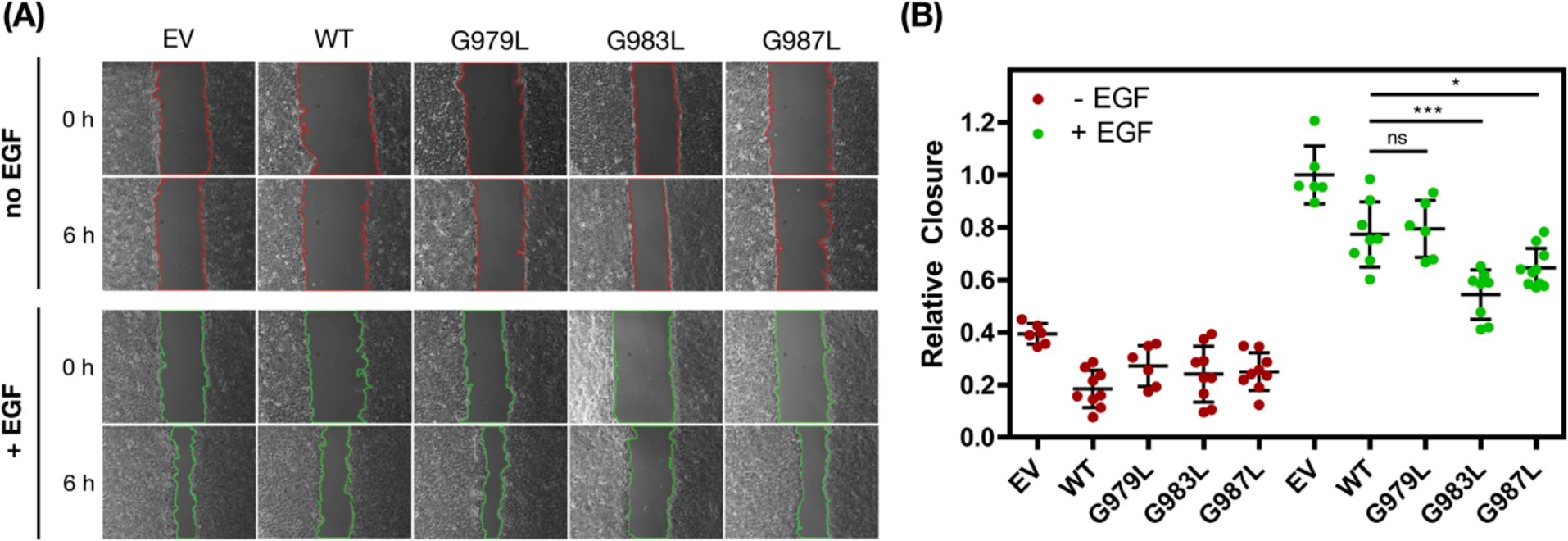
PTPRJ point mutants inhibit EGFR-driven cell migration. Representative phase contrast images with tracings to identify open scratch areas **(A)** and quantification **(B)** of the effect of PTPRJ point mutations on wound closure. UMSCC2 were serum-starved, scratched (0 h) and incubated with serum-free media with or without EGF (50 ng/mL) for 6 h. Relative closure was quantified by calculating the percent change in area between 0 and 6 h using ImageJ, and then normalized to EV (+ EGF). **(B)** Results are shown as mean ± SEM (*n* = 6-9). Statistical significance between WT and mutants was assessed using one-way ANOVA corrected for multiple comparison (Dunnet test) at 95% confidence intervals): \*\*\**p* ≤ 0.001; and \**p ≤* 0.05

### PTPRJ point mutant disrupts full-length PTPRJ oligomerization and promotes its access to EGFR

To determine whether changes in phosphorylation and cell migration were direct functional consequences of the disruption of PTPRJ homodimers, we assessed the oligomerization of full-length PTPRJ WT and G983L (i.e., the mutant disrupting self-association, kinase phosphorylation, and cell migration the most) expressed in UMSCC2 cells by *in situ* proximity ligation assay (29-31). Figure 4 shows that strong PLA signal was observed with WT PTPRJ, indicating that PTPRJ exists as oligomers. Strikingly, a significant decrease in PLA signal was observed in cells expressing the receptor with the G983L point mutant compared to those with WT, confirming that the G983L point mutation disrupts PTPRJ oligomerization, and significantly reduces the number of PTPRJ oligomers.

**Figure 4.**
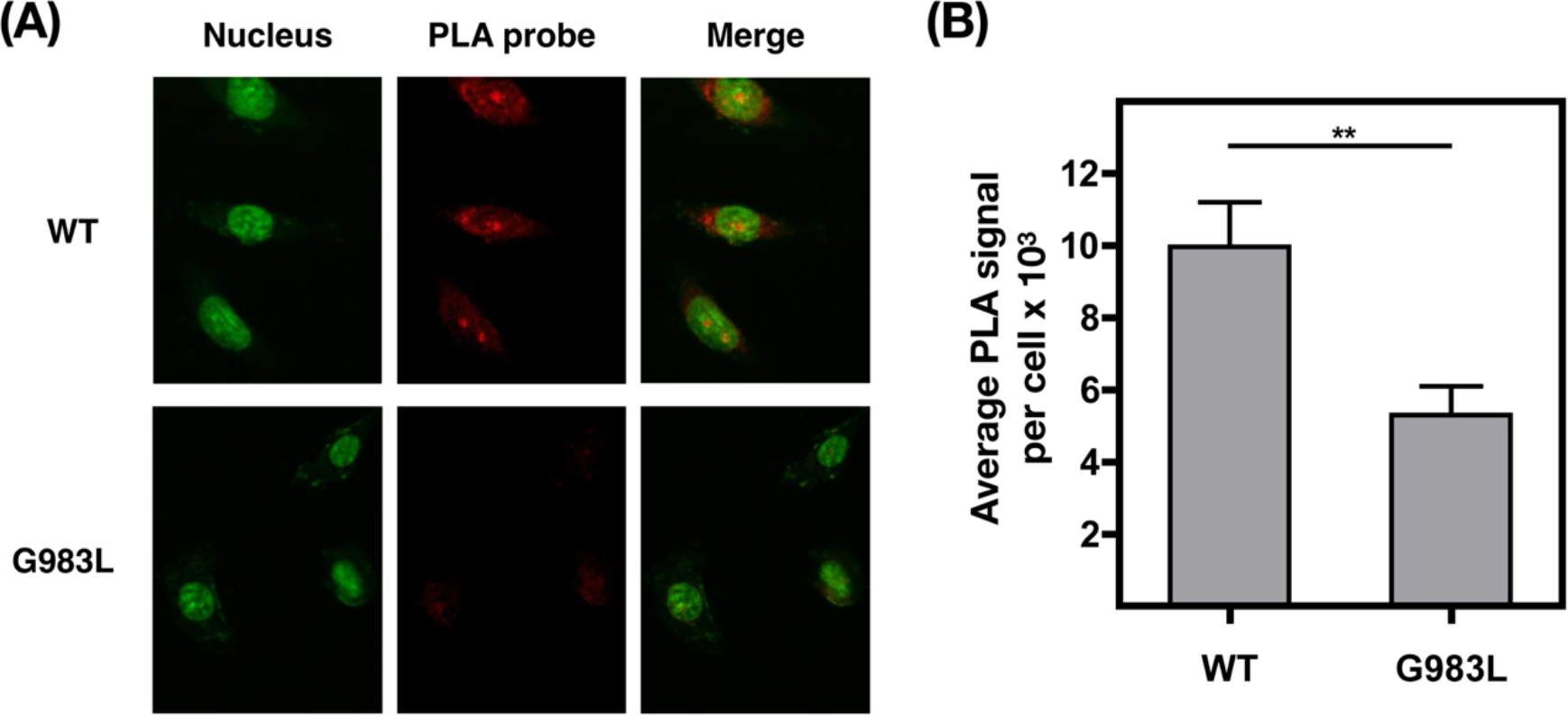
Full-length PTPRJ point mutant disrupts PTPRJ oligomerization in cells. Representative confocal microscopy images **(A)** and quantification **(B)** of the effect of transmembrane point mutation on PTPRJ oligomerization using *in situ* PLA. Oligomerization of full-length PTPRJ WT and G983L expressed in UMSCC2 cells was measured by spinning disk confocal microscopy (green: nucleus; red: PLA for PTPRJ self-association). The average PLA signal intensity per cell was determined using BlobFinder. **(B)** Results are shown as mean ± SEM (*n* > 100). Statistical significance was assessed using unpaired *t* test (at 95% confidence intervals): \*\**p* ≤ 0.01

Moreover, since physical interaction of PTPRJ with RTKs, such as EGFR, appears integral to its regulatory role of RTK signaling (10, 29), we investigated whether destabilizing full-length PTPRJ oligomerization promotes its interaction with EGFR. Figure 5 shows that cells expressing the G983L PTPRJ point mutant exhibited a stronger PLA signal between PTPRJ and EGFR than cells expressing wildtype PTPRJ. This suggests that, as hypothesized, disrupting PTPRJ self-association favors its access to EGFR.

**Figure 5.**
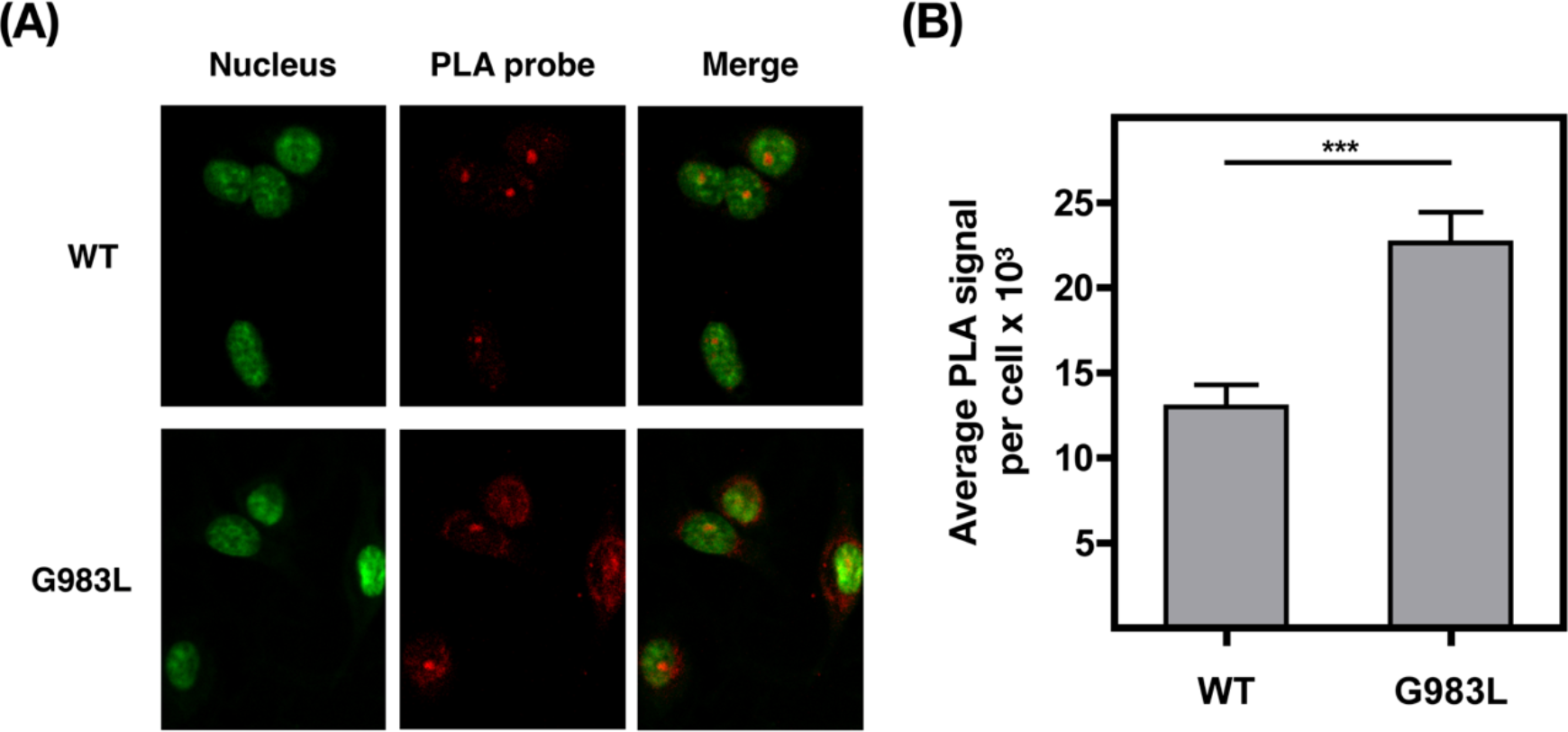
Full-length PTPRJ point mutant promotes interaction with EGFR in cells. Representative confocal microscopy images **(A)** and quantification **(B)** of the effect of transmembrane point mutation on PTPRJ interaction with EGFR using *in situ* PLA. Complex formation with EGFR was measured by spinning disk confocal microscopy in UMSCC2 cells expressing WT or G983L PTPRJ (green: nucleus; red: PLA between PTPRJ and EGFR). The average PLA signal intensity per cells was determined using BlobFinder. **(B)** Results are shown as mean ± SEM (*n* > 100). Statistical significance was assessed using unpaired *t* test (at 95% confidence intervals): \*\*\**p* 0.001

Altogether, our results indicate that TM domain association stabilizes PTPRJ oligomerization through a specific contact interface consisting of G979, G983 and G987 (i.e., GxxxGxxxG motif) and that disrupting PTPRJ self-association through point mutations promotes PTPRJ phosphatase activity, likely by favoring its access to EGFR. Our results also suggest that this structure-function relationship may provide a new opportunity to augment PTPRJ activity in cancer cells driven by EGFR signaling.

### Design and identification of a TM peptide sequence capable of antagonizing PTPRJ self-association

To leverage the identified regulatory TM domain interactions of PTPRJ, we set out to develop exogenous peptides that can modulate PTPRJ enzymatic activity by disrupting receptor oligomerization. To do so, we constructed a library of peptides designed to selectively bind to the TM domain of PTPRJ (see *Experimental procedures*). Similar peptide library design approaches have proven successful at identifying and selecting peptides to inhibit viral TM domain dimerization as well as transmembrane receptor dimerization (32, 33). The degenerative oligonucleotide sequence corresponding to the designed TM binder consensus peptide sequence (Figure 6A) was subcloned in fusion with AraC* to be tested against WT PTPRJ-AraC, using the DN-AraTM assay (34). After co-transforming these constructs into *E. coli*, individual colonies were tested for GFP emission. Of the 299 individual colonies that were screened (Figure 6B), 28 clones (Figure 6B, green dots) showed greater disruption of PTPRJ self-association than the wild-type PTPRJ construct (Figure 6B, red line). After further DN-Ara-TM validation and DNA sequencing, the sequence of the clone that consistently showed the most GFP signal disruption was selected as the lead peptide sequence (Figure 6C, lead). Importantly, this lead peptide sequence showed greater disruption of PTPRJ self-association than the WT sequence as designed (Figure 6D). Next, we set out to identify residues important for the interaction of this lead peptide binder sequence with the TM domain of PTPRJ. Guided by preliminary molecular simulations indicating that Ser7, Ala11 and V15 (Figure 6C) may be part of the contact interface with PTPRJ (data not shown), we mutated these 3 residues to leucines with the goal to disrupt helix packing, and tested their effect on PTPRJ self-association using the DN-AraTM assay. Figure 6E shows that all three point-mutations significantly hinder the ability of the lead peptide sequence to disrupt PTPRJ self-association, with A11L having the most effect.

**Figure 6.**
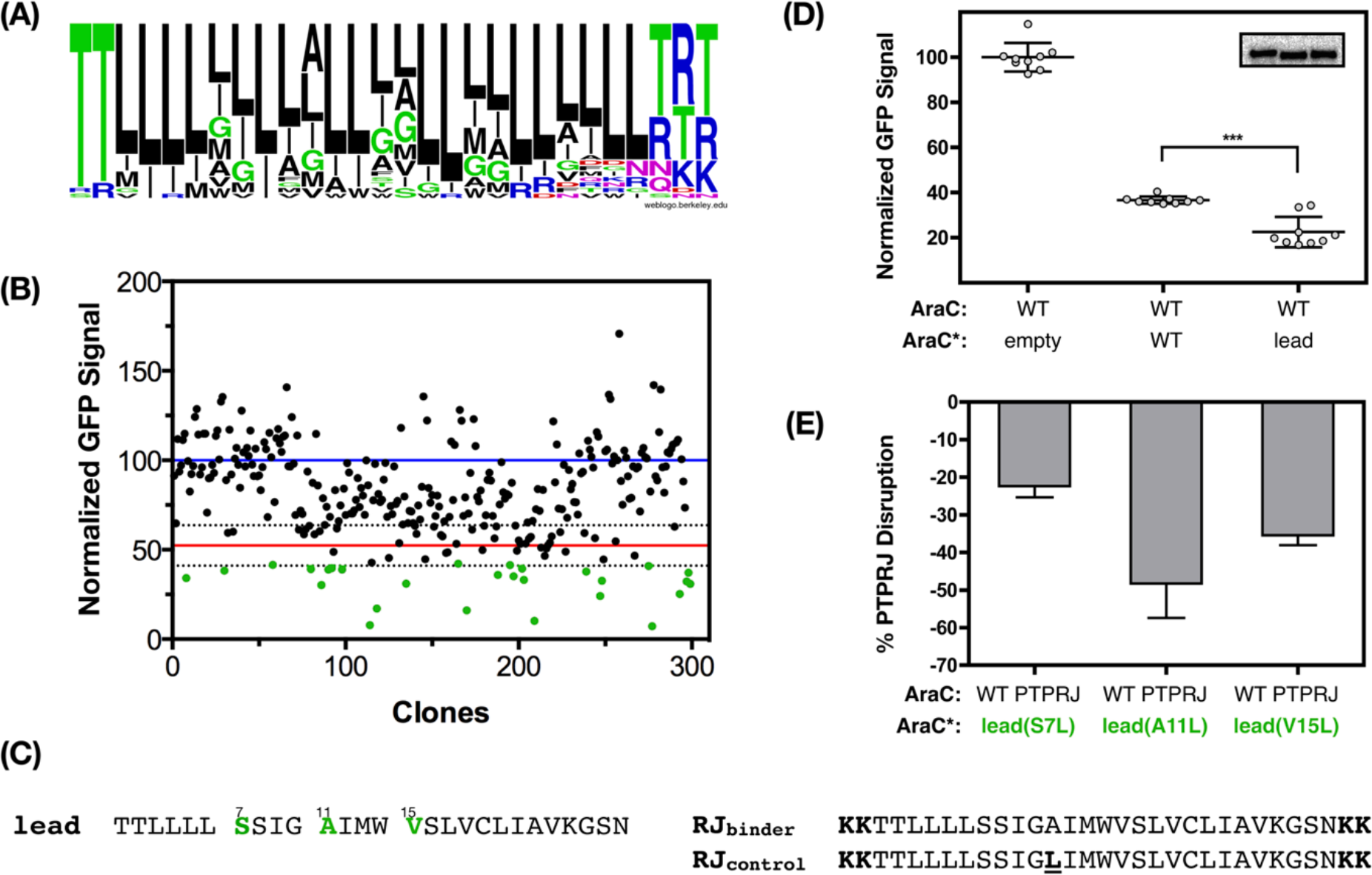
Design and identification of a peptide binder against the TM domain of PTPRJ. **(A)** Consensus peptide sequence of TM binder against PTPRJ designed. (**B)** Distribution of AraTM results of 299 TM constructs cloned in fusion with AraC* in competition against the WT PTPRJ sequence. Average signals from the PTPRJ dimer and from the competition with the WT sequence are represented as a blue and red line, respectively. Dotted lines represent one standard deviation. Green dots represent top hits that were sequenced and retested. **(C)** Amino acid sequence of the identified and selected lead, and the sequences of the synthesized RJ_binder_ and RJ_control_ peptides. **(D)** Assessing the disruption of PTPRJ oligomerization by the identified lead peptide binder using the DN-AraTM assay. Results are normalized to PTPRJ WT signal and are shown as mean ± SEM (*n* = 9). Statistical significance was assessed using unpaired *t* test (at 95% confidence intervals): \*\*\**p* ≤0.001. **(D, inset)** Representative immunoblot against MBP. **(E)** Identifying residues important for the interaction of the lead sequence with the TM domain of PTPRJ. DN-AraTM results of lead point mutants in competition against WT PTPRJ. Results are normalized to the wild type lead and shown as mean ± SEM (*n* = 9). (**E, inset)** Representative immunoblot against MBP, showing similar expression level

### Interaction of RJ_binder_ with Membrane Mimics and Cell Membranes

Peptides corresponding to the lead and the control A11L sequences (referred to subsequently as RJbinder and RJ_control_, respectively) were synthesized and purified (Figure S4). Due to the relatively high hydrophobicity of the peptide sequences, two lysine residues were appended to the N- and C-termini to enhance water solubility and facilitate purification (Figure 6C) (35). However, despite these modifications, solubilization of such peptides in detergent is required to prevent aggregation in solution and allow partitioning into cell membranes (36-39). Based on the design approach and the hydrophobicity of these peptides, one would expect that they insert and fold as TM α-helices in both detergent and lipid membrane mimics. Circular dichroism spectroscopy shows that both RJ_binder_ and RJ_control_ fold into α-helices in the presence of n-octylglucoside micelles (39) and POPC large unilamellar vesicles (Figure S4 A and B). Proper membrane insertion in both membrane mimetic environments was also confirmed by fluorescence emission measurements: A significant blue shift in tryptophan fluorescence emission (> 10 nm) was observed when peptides were incubated with either n-octylglucoside micelles or POPC large unilamellar vesicles (Figure 6C and D), blue line) when compared to fluorescence emission in buffer. Taken together, these results indicate that both RJ_binder_ and RJ_control_ adopt a α-helical transmembrane conformation in membrane mimetic environments. We also explored whether RJ_binder_ can incorporate into UMSCC2 cell membranes. To this end, we studied the cell membrane partition of a fluorescently labeled version of RJ_binder_ initially incorporated into n-octylglucoside micelles. To allow peptide partition into cell membranes, peptide/octylglucoside solutions were added to cells to obtain a final detergent concentration (3 mM) well below its critical micellar concentration (~ 15 mM as measured in cell media). Confocal microscopy showed the fluorescently-labeled RJ_binder_ peptide to be localized on the cell surface (Figure S5).

### RJ_binder_ disrupts PTPRJ self-association and inhibits EGFR signaling

Next, we investigated the effect of exogenously adding RJ_binder_ to UMSCC2 cells on PTPRJ dimerization and cell phenotypes. We hypothesized that, similarly to dimerization-disrupting point mutants (e.g., G983L), treating cells with RJ_binder_ (initially incorporated into n-octylglucoside micelles) would inhibit not only receptor dimerization, but also EGFR signaling. Figure 7 shows that a significant decrease in PLA fluorescence average signal was observed following treatment of cells expressing WT PTPRJ with RJ_binder_, indicating the ability of the peptide to disrupt PTPRJ self-association. Moreover, significant inhibition of EGFR phosphorylation (at both Y1068 and Y1173) (Figure 8), and cell migration was observed in cells treated with RJ_binder_ (Figure 9). We also hypothesized that RJ_binder_ should have little effect (if any) on cells expressing either minimal PTPRJ (EV) or the homodimer-disrupting mutant G983L (as the PTPRJ homodimer is already destabilized). Figure S6 shows that RJ_binder_ indeed had no effect on cell migration when added to these two cell lines. Crucially, cell treatment with RJ_control_ did not affect receptor self-association (Figure 7), EGFR phosphorylation (Figure 8) or wound closure (Figure 9). Altogether, these results imply that the specific inhibitory effects observed in cells were due to a direct interaction of RJ_binder_ with the cell endogenous PTPRJ receptor.

**Figure 7.**
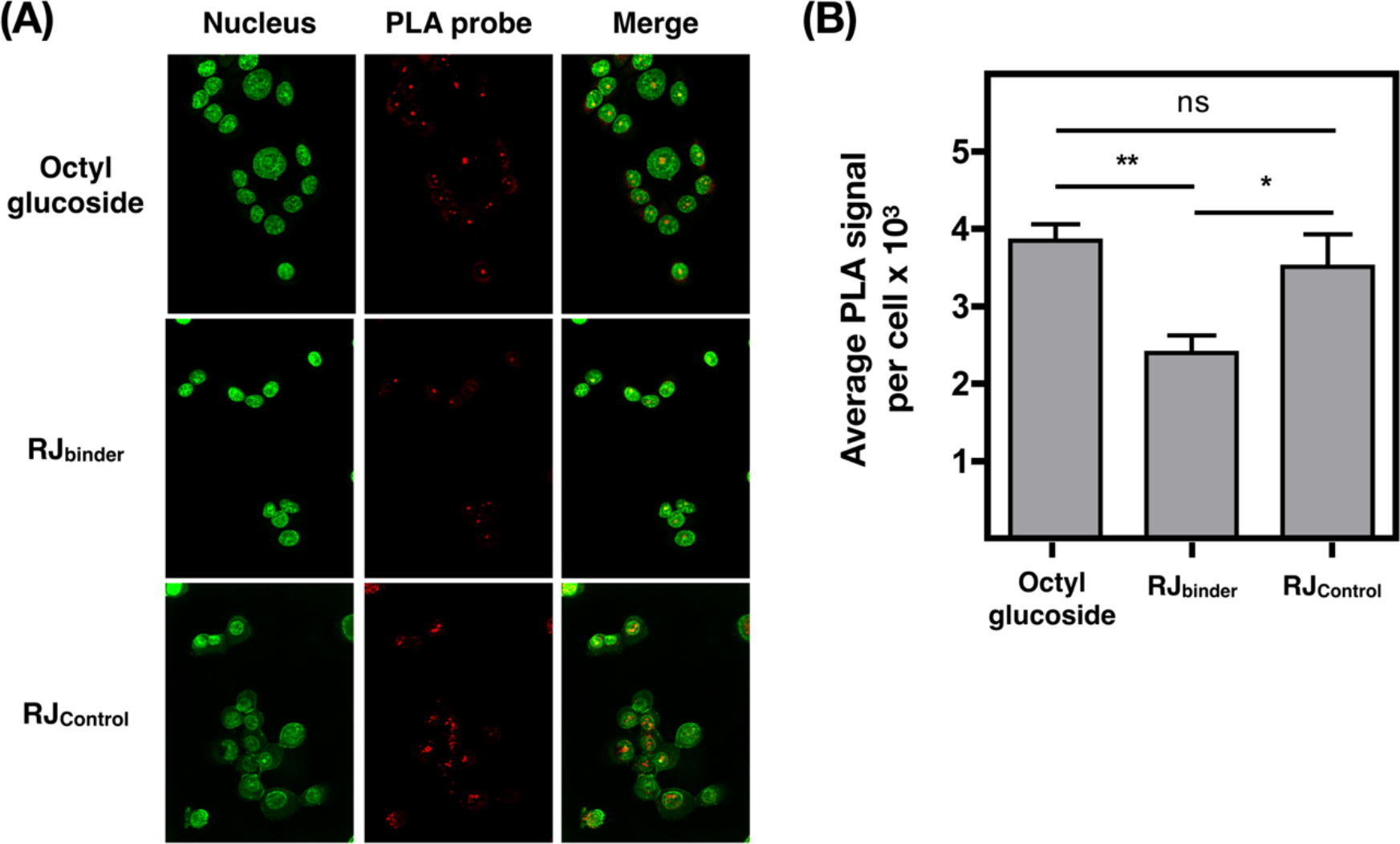
RJ_binder_ disrupts full-length PTPRJ self-association in cells. Representative confocal microscopy images **(A)** and quantification **(B)** of the effect of RJ_binder_ and RJ_control_ on PTPRJ oligomerization using *in situ* PLA. Oligomerization of full-length PTPRJ WT in UMSCC2 cells was measured following treatment with RJ_binder_ or RJ_control_ for 1 h using spinning disk confocal microscopy (green: nucleus; red: PLA). The average PLA signal intensity per cells was determined using BlobFinder. **(B)** Results are shown as mean ± SEM (*n* > 100). Statistical significance was assessed using one-way ANOVA corrected for multiple comparison (Dunnet test) at 95% confidence intervals): \*\**p* ≤ 0.01 and \**p* ≤ 0.05

**Figure 8.**
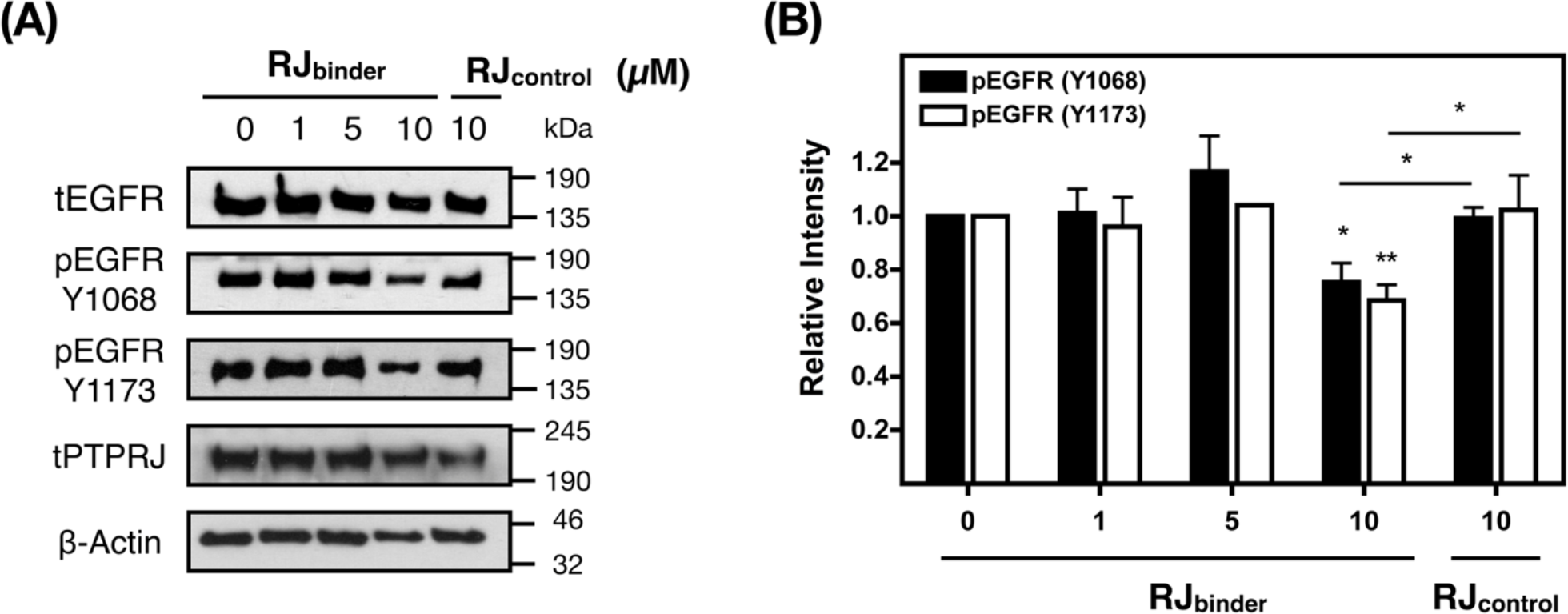
RJ_binder_ inhibits the phosphorylation of EGFR. **(A)** Serum-starved UMSCC2 cells were treated with RJ_binder_ or RJ_control_ for 3 h prior to stimulation with EGF. Cell lysates were probed for EGFR, phospho-EGFR (Y1068 and Y1173), and PTPRJ by immunoblot. **(B)** The relative (ratio of phosphorylated to total protein) intensities are shown as mean ± SEM (*n* = 3-5). Statistical significance was assessed using unpaired *t* test (at 95% confidence intervals): \*\**p* ≤ 0.01 and \**p* ≤ 0.05

**Figure 9.**
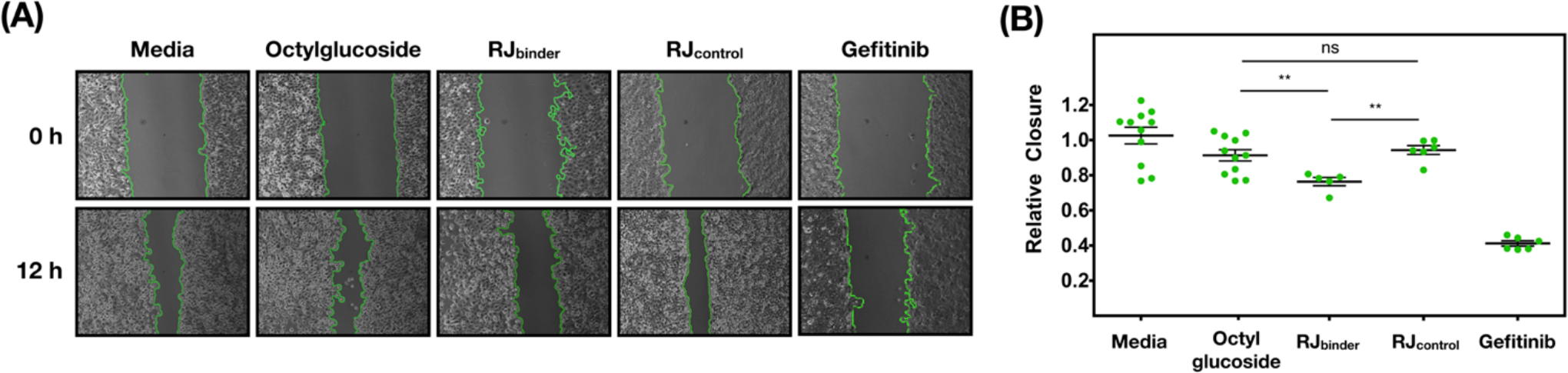
RJ_binder_ inhibits EGFR-driven cell migration. Representative phase contrast images with tracings to identify open scratch areas (A) and quantification **(B)** of the effect of RJ_binder_, RJ_control_, and gefitinib on wound closure. Serum-starved UMSCC2 cells were treated with 3 mM n-octylglucoside, 10 μM RJ_binder_, 10 μM RJ_control_, or 15 μM gefitinib (a known inhibitor of EGFR activity) for 1 h, scratched (0 h), and incubated in media with EGF (50 ng/mL) for 12 h. Relative closure was quantified by calculating the percent change in area between 0 and 12 h using ImageJ, and then normalized to media containing EGF. **(B)** Results are shown as mean ± SD (*n* = 6-9). Statistical significance between WT and mutants was assessed using one-way ANOVA corrected for multiple comparison (Dunnet test) at 95% confidence intervals): \*\*\**p* ≤ 0.001; and \**p ≤* 0.05

## Discussion

The main goal of this work was to extend our structure-function understanding of the role of TM-mediated interactions in modulating the activity of PTPRJ. We showed that PTPRJ dimerization is mediated by a specific single-contact TM interface constituted by two overlapping GxxxG motifs, a common motif for TM-TM homodimerization. Interestingly, when the corresponding full-length PTPRJ glycine point mutants (G979L, G983L and G987L) were individually expressed in UMSCC2 cells, we observed a decrease in EGFR and ERK phosphorylation, and inhibition of EGFR-driven wound closure. We also showed that expression of the most disruptive PTPRJ mutant (G983L) led to a decrease in full-length PTPRJ homodimer formation in cells when compared to the wild type receptor, validating the importance of this particular glycine residue in mediating receptor dimerization. As anticipated, the suppression of phosphorylation, wound closure and dimerization, while significant, is not complete. There are several potential reasons: interfering with a single amino acid is unlikely to completely disrupt full-length receptor self-association in cell membranes; TM-TM interaction may not be the only mediator of PTPRJ dimerization (1, 40); and EGFR is known to be regulated by a number of other PTPs, such as PTPRS, PTPRK, LAR, SHP-1 and PTP1B (41).

Interestingly, we also observed an increase in PLA signal between PTPRJ and EGFR in cells expressing the G983L PTPRJ point mutant, suggesting that destabilizing PTPRJ self-association promotes its access to EGFR. While proximity detected by *in situ* PLA does not provide evidence of direct EGFR-PTPRJ interaction, the observed enhanced association, in addition to the decreased EGFR phosphorylation, strongly argues for a specific and functional interaction. It is then compatible with our hypothesis that PTPRJ TM-mediated dimerization prevents access to its substrates. Furthermore, these results and interpretations are consistent with the physical interaction at the cell surface between EGFR and PTPRJ identified by Yarden and coworkers (10). The same study reported that, while EGFR undergoes endocytosis after interacting with PTPRJ at the plasma membrane, PTPRJ remains confined to the cell surface. It is then reasonable to hypothesize that EGFR endocytosis would be forestalled by enhanced dephosphorylation by and/or association with G983L PTPRJ. These effects on EGFR internalization are the focus of on-going investigations in our laboratories.

In the present study, with the overarching goal to develop novel PTPRJ agonist peptides, we also rationally designed and selected a peptide (RJ_binder_) capable of binding to the TM domain of PTPRJ and antagonizing its self-association. Remarkably, treating UMSCC2 cells expressing WT PTPRJ with RJ_binder_ led to inhibition of phosphorylation, wound closure and PTPRJ self-association. Similar to the effect of TM point mutations, RJ_binder_ 10 μM treatments did not completely suppress EGFR phosphorylation. It could be due to the same reasons discussed above for the effect of single-point mutations and for the fact that interfering with TM-TM interaction may not be sufficient to completely overcome receptor self-association (39). Because peptides are reconstituted in detergent micelles prior to incubation with cells, another explanation is that the population of inserted peptides is a mixture of both possible orientation (i.e., N-terminus or C-terminus out), which would contribute to a lack of efficacy against PTPRJ. It is also conceivable that the activity of RJ_binder_ may be highly cell-context dependent as possible RTK substrates and PTPRJ itself are likely to be differentially expressed across cancer cell type. Its activity could also be affected by the high level of PTPRJ expression observed in the transfected UMSCC2 cells, against which the peptide has to compete. It is therefore informative to compare the activity of the RJ_binder_ with gefitinib, a potent inhibitor of EGFR activity used in cancer treatment. Indeed, cell treatment with 15 μM of gefitinib (IC_50_ for UMSCC2 cells expressing WT PTPRJ; Figure S7), suppresses wound closure by ‘only’ ~60% (Figure 9), showing the importance of the targeted cells. Moreover, RJ_binder_ showed no activity against cells expressing either minimal level of PTPRJ or the homodimer-disrupting mutant G983L, and the control peptide with hindered interaction with PTPRJ (i.e., RJ_control_) showed no effect on receptor self-association, phosphorylation or wound closure. Altogether, these results indicate that RJ_binder_ interacts specifically with the TM domain of PTPRJ.

It is also important to point out that RJ_binder_ represents a proof-of-concept for a new allosteric and possibly orthogonal way to target the activity of RPTPs and receptor kinase phosphorylation. Indeed, while other methods have been devised to modulate RPTPs (e.g., small molecules targeting the phosphatase domain, wedge peptide mimetic and decoy protein), most, if not all, act by inhibiting their activity and none of them target the TM domains (14). Moreover, the mechanism of activation of RPTPs remain unclear and no common model has been proposed. As mentioned in the introduction, it is due, in part, to the lack of known natural ligands that are able to trigger RPTPs intracellular pathways by binding to the extracellular domain in the same manner than with RTKs, for example. For instance, only two biological ligands have been identified for PTPRJ: heparan sulfate proteoglycan syndecan-2 (42) and thrombospondin-1 (43), both of which are able to bind to the extracellular domain and stimulate PTPRJ activity. However, there is no structural model for their mechanism of action. Therefore, being able to activate PTPRJ allosterically using TM peptide binders can provide significant insight into how this receptor functions, interacts, and eventually be modulated.

Because of its tumor-suppressor role, there are clear advantages in promoting PTPRJ activity towards not only EGFR, but also other deregulated RTKs in cancer cells. While fully suppressing EGFR phosphorylation through disruption of PTPRJ TM-TM interaction remains to be explored, PTPRJ is known to suppress other growth factor receptors and their signaling proteins, including VEGF receptor 2 (44, 45), hepatocyte growth factor receptor (46-48), FGF receptor (46), and PGDF receptor (PDGFR) (45), and the RAS- and FLT3-mediated signaling pathways by dephosphorylating ERK1/2 (29, 49). Therefore, disrupting PTPRJ self-association may inhibit other RTKs known to be involved in cancer cell progression, which may potentiate the effect of a peptide such as RJ_binder_.

It is also true that activating PTPRJ may have potential unintended consequences on cell survival. For instance, in addition to these antiproliferative and tumor suppressor functions, PTPRJ activity may be driving oncogenic signaling through positive regulation of VEGF-induced Src and Akt activation, as well as of endothelial cell survival (45). However, one would expect that the role of PTPRJ, the consequences of inhibiting/activating it, and the magnitude of these effects to be cell type-dependent. Improving the efficacy of RJ_binder_ and quantitatively determining the cancer cell context-dependent outcomes of interfering with PTPRJ activity through TM mutations and TM-directed peptide binders is the focus of on-going efforts in our laboratories. Moreover, since RJ_binder_ appears to act allosterically and by activating PTPRJ, it is tempting to speculate that such peptides could potentiate the activity of conventional small molecule inhibitors that often also have incomplete suppression of downstream signaling pathways or overcome acquired drug resistance when secondary mutations alter small molecule binding. Finally, we expect that the basic framework developed here can be extended to other RPTPs because modulation of RPTP activity through dimerization of TM domains appears to be a general theme throughout the family, and TM domains may provide orthogonality not possible by targeting either the extracellular or PTP domains. Therefore, this approach may afford a new modality to study RPTPs activity and represent a novel class of therapeutics against RTK-driven cancers.

## Methods

### Subcloning

Unless otherwise stated, standard molecular biology techniques were used, and all constructs used were verified by DNA sequencing (Genewiz, Inc.). The DNA sequence coding for the TM domains of interest were cloned into either pAraTMwt (coding for AraC) or pAraTMDN (coding for the inactive form of AraC, AraC*) plasmids. The protein sequence of human PTPRJ used in AraTM assays contains five extracellular residues, the TM domain, and twenty cytoplasmic juxtamembrane residues:

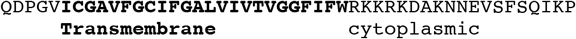

The sequence coding for full-length human PTPRJ (Entrez Gene ID 5795; PlasmID Repository) was subcloned into the pBABE-puro retroviral vector as an EcoRI/SalI fragment. All site-directed mutagenesis was performed using the QuikChange II Site-Directed Mutagenesis Kit (Agilent).

### DN-AraTM Dimerization Assay

The constructs and the reporter plasmid (pAraGFPCDF) were co-transformed into the AraC-deficient *E. coli* strain SB1676 and streaked onto selective plates. Colonies were picked from each construct and grown in LB media for 8 h at 30 °C. Each culture was diluted into selective auto-induction media and grown for an additional 16 h at 30 °C. A series of 2-fold dilutions of the cultures were prepared in a black 96-well, clear-bottom plate. Absorption at 580 (10) nm and GFP fluorescence emission spectra (excitation maximum 485 (20) nm and emission maximum at 530 (30) nm were collected using an Infinite^®^ 200 PRO Plate Reader (Tecan). The results are reported as the ratio of fluorescence emission at 530 nm to absorbance at 580 nm and normalized to the negative control (empty plasmids and reporter plasmid). Immunoblotting was performed using HRP-conjugated anti-maltose binding protein (MBP) monoclonal antibody at 1:10000 dilution (New England Biolabs, #E8038) to verify equal expression levels of each construct.

### Maltose Complementation Test

Plasmids pAraTMwt and pAraTMDN containing TM inserts were transformed into the MBP-deficient *E. coli* strain MM39 and streaked onto selective lysogeny broth (LB) plates. The following day, individual colonies were picked and grown in LB media. Saturated culture from each construct was streaked onto selective M9 minimal media plates containing 0.4% (w/v) maltose and incubated for three days at 37°C.

### Cell Treatment for Immunoblot Analyses

Human squamous cell carcinoma UMSCC2 cells (gift of Prof. Alexander Sorkin, University of Pittsburgh) were cultured in Dulbecco’ Modified Eagle’ medium (DMEM) high glucose supplemented with 10% FBS, 100 units/ml penicillin and 0.1 mg/ml streptomycin. Cells were cultured in a humidified atmosphere of 5% CO_2_ at 37 °C. To express full-length PTPRJ in UMSCC2 cells, pBABE-puro retroviral vector constructs were first transduced into retrovirus-producing Phoenix cells (Dr. Gary Nolan, Stanford University) using the calcium phosphate-DNA co-precipitation method in the presence of chloroquine. About 60 h post transfection, viral particles were collected, passed through a 0.45 μM filter and added to UMSCC2 cells. Positive transductants were selected with puromycin.

### Cell Culture and Stable Transduction

UMSCC2 cells were seeded in 12 well plates at 400,000 cells/well and incubated overnight. The cell media was replaced with serum-free media for 2 h before treatment. For peptide treatments, RJ_binder_ and RJ_control_ were resuspended in 30 mM n-octylglucoside to obtain a 100 μM final peptide solution. Peptide/octylglucoside solutions were added to serum-starved cells to obtain a final detergent concentration (3 mM) lower than its critical micellar concentration to allow peptide partition into cell membrane (39). Following a 3 h treatment at 37 °C, treatment media was removed and replaced with fresh serum-free media containing EGF (10 ng/mL), and the cells were incubated for another 10 min at 37 °C. Cells were solubilized by the addition of cell extraction buffer supplemented with broad-spectrum phosphatase and protease inhibitors (Pierce #88667), then removed and analyzed by immunoblot.

### Immunoblot Analysis

Samples were boiled for 10 min at 95 °C and resolved by SDS-PAGE on an 8% tris-glycine gel. Subsequently, the samples were transferred onto a 0.45 μm nitrocellulose membrane (GE Healthcare #1060002) at 100 V for 1 h at 4 °C. Membranes were blocked with 5% bovine serum albumin (BSA) in tris-buffered saline Tween 20 (TBS-T) for 1 h at RT and then blotted for phosphorylated proteins (Cell Signaling Technology; phospho-EGFR Y1068 #3777, phospho-EGFR Y1173 #4407, phospho-p44/42 MAPK (ERK1/2) #4377). When blotting for total proteins (Cell Signaling Technology; EGFR #4267, p44/42 MAPK #4695, β-Actin #3700 and Santa Cruz Biotechnology; PTPRJ #376794), the membranes were blocked with 5% dry milk in TBS-T for 1 h at RT. Following blocking, the membranes were incubated with primary antibodies in 5% BSA TBS overnight at 4 °C (pEGFR at 1:1000 dilution, total EGFR 1:2000, PTPRJ, 1:100, β-Actin 1:4000). Membranes were then incubated with the appropriate secondary antibodies in TBS-T for 30 min at RT at 1:4000 dilution (Cell Signaling Technology, Anti-rabbit #7074, Anti-mouse #7076). Following washes with TBS-T, the immunoblot was visualized by chemiluminescence after incubation with Clarity Western ECL Substrate (Bio-Rad). Images were quantified using ImageJ and plotted as normalized (ratio of phosphorylated to total intensity) mean values.

### Scratch Assay

UMSCC2 cells were seeded in 6-well plates at a cell density necessary to reach confluency after 24 h. Cells were incubated with serum-free media for 2 h, after which a scratch was made in the confluent cell monolayer with a 200 μL pipette tip to create a thin gap. Cells were immediately washed once with serum-free media and imaged using phase contrast microscopy (*t* = 0 h). The cells were incubated with serum-free media containing EGF (50 ng/mL) to stimulate scratch closure or medium lacking EGF (control) at 37 °C for 6 h at which point phase contrast images were taken (*t* = 6 h) with a 10x objective using an Eclipse Ti-S microscope. For peptide treatments, RJ_binder_ and RJ_control_ were prepared as previously described for immunoblot analyses and incubated at 37 °C for 1 h before the initial scratch. The media was then replaced with complete media containing EGF (50 ng/mL) at 37 °C for 12 h. Scratch areas were quantified with ImageJ using the MRI Wound Healing Tool, and the closure percent was found by calculating the percent change in area between the initial and final time points. Percent scratch closure was normalized to either empty pBabe-puro vector (EV) or wild-type cells treated with media containing EGF (50 ng/mL).

### Proximity Ligation Assay

UMSCC2 cells were seeded on 22 mm × 22 mm coverslips in 6 well plates at 300,000 cells/well to be ~60% confluent after 16 h. For peptide treatments, RJ_binder_ and RJ_control_ were prepared as previously described for immunoblot analyses and incubated with the cells in complete media at 37 °C for 1 h. Following 3-5 washes with PBS, the cells were fixed and permeabilized with ice cold methanol for 10 min at RT. The coverslips were blocked in 1% BSA in PBS at RT for 30 m and washed 3 times with PBS. The coverslips were incubated overnight in a humidified chamber at 4 °C with primary antibodies (anti-PTPRJ mouse and anti-PTPRJ rabbit at 1:50 dilutions). Following the manufacturer protocol (Sigma-Aldrich; Duolink^®^ #DUO92101), the coverslips were then washed with PBS. The PLUS and MINUS PLA probes were diluted in the Duolink^®^ Antibody Diluent (1:5), and the coverslips were incubated in a humidity chamber for 1 h at 37 °C. After washing with the supplied Buffer A, the coverslips were incubated with the provided ligase in a humidity chamber for 30 min at 37 °C. After a second wash with Buffer A, the cells were incubated with the provided polymerase in amplification buffer in a humidity chamber for 100 min at 37 °C. The coverslips were then washed stained with Nucspot^®^ Live 488 (Biotium #40081) and mounted on slides with Fluoromount (Sigma-Aldrich #F4680) before being imaged by spinning disc confocal microscopy. The images were analyzed and quantified using BlobFinder (Center for Image Analysis).

### Peptide TM Binder Library Construction

To design a library of peptide sequences to be screened for their ability to compete for PTPRJ self-association using DN-AraTM, we first generated sequence alignments based on the wild-type PTPRJ TM sequence. From this alignment, variability at each aligned position within the TM sequence was determined; positions where more than different 4 amino acids occurred within the alignment were chosen for the library, excluding positions where conservative substitutions only occurred. We also retained key amino acids important for dimer formation, particularly the small-x3-small motif present in PTPRJ and focused on varying positions flanking these key residues; previous approaches have shown that residues flanking core small-x3-small motif residues can modulate specificity of dimerization. Based on these results, a consensus peptide sequence was generated (Figure 6) and a corresponding degenerate codon sequence was chosen to capture as much of the sequence diversity as possible while minimizing the number of different degenerate codons required. Similar degenerate codon sequences and approaches have been used previously in library design and selection of TM peptides (32, 33).

### Solid-phase Peptide Synthesis

RJ_binder_ (H_2_N-KKTTLLLSSIGAIMWVSLVCLIAVKGSNKK-CONH_2_) and RJ_control_ (H_2_N-KKTTLLLSSIGLIMWVSLVCLIAVKGSNKK-CONH_2_) peptides were prepared by Fmoc solid-phase chemistry using a CEM Liberty Blue microwave synthesizer. For fluorescence microscopy experiments, 5(6)-carboxyfluorescein was conjugated to the N-terminus of RJ_binder_ (FITC-RJ_binder_) on resin using standard coupling protocol. Briefly, peptides were purified via reverse-phase high-performance liquid chromatography (RP-HPLC; Phenomenex Luna Omega 5 μm 250 × 21.2 mm C18; flow rate 5 mL/min; phase A: water 0.1% TFA; phase B: acetonitrile 0.1% TFA; gradient 60 min from 95/5 A/B to 0/100 A/B). The purity of the peptides was determined by RP-HPLC (Phenomenex measurements in vesicles, RJ_binder_ and RJ_control_ peptides were solubilized with hexafluoro-2-propanol and incubated with 1-palmitoyl-2-oleoyl-sn-glycero-3-phosphocholine (POPC) at a 1:300 ratio. The peptide and POPC mixture was dried as a thin film and held under vacuum for at least 24 h. The mixture was rehydrated in 10 mM HEPES, 10 mM KCl, pH 8.0, for at least 30 min with periodic gentle vortexing. The resulting large multilamellar vesicles containing RJ_binder_ or RJ_control_ were freeze-thawed for seven cycles and subsequently extruded through a polycarbonate membrane with 100 nm pores using a Mini-Extruder (Avanti Polar Lipids) to produce large unilamellar vesicles.

### CD Spectroscopy

Far-UV CD spectra were recorded on a Jasco J-815 CD spectrometer equipped with a Peltier thermal-controlled cuvette holder (Jasco). Measurements were performed in 0.1 cm quartz cuvette. CD intensities are expressed in mean residue molar ellipticity [θ] calculated from the following equation:

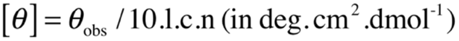

where, *θ_obs_* is the observed ellipticity in millidegrees, *l* is the optical path length in centimeters, *c* is the final molar concentration of the peptides, and *n* is the number of amino acid residues. Raw data was acquired from 260 to 200 nm at 1 nm intervals with a 100 nm/min scan rate, and at least five scans were averaged for each sample. The spectrum of Luna Omega 5 μm 250 × 10 mm C18; flow rate 5 mL/min; phase A: water 0.1% TFA; phase B: acetonitrile 0.1% TFA; gradient 60 min from 95/5 A/B to 0/100 A/B), and their identity was confirmed via matrix-assisted laser desorption/ionization time of flight (MALDI-TOF) mass spectroscopy (Figure S3).

### Sample Preparation of CD and Tryptophan Fluorescence Measurements

Lyophilized RJ_binder_ and RJ_control_ were solubilized to 20 μM in 10 mM HEPES, 10 mM KCl, pH 8.0, and incubated for 1 h at RT. Each construct was diluted to a final concentration of 10 μM before analysis. For measurements in micelles, lyophilized RJ_binder_ and was solubilized to 10 μM with 30 mM n-octylglucoside prepared in 10 mM HEPES, 10 mM KCl, pH 8.0. For octylglucoside micelles or POPC liposomes was subtracted out from all construct samples.

### Tryptophan Fluorescence Spectroscopy

Fluorescence emission spectra were acquired with a Fluorolog-3 Spectrofluorometer (HORIBA). The excitation wavelength was set at 280 nm and the emission spectrum was measured from 300 to 450 nm. The excitation and emission slits were both set to 5 nm.

### Peptide Incorporation into Cell Membrane by Fluorescence Microscopy

RJ_binder_ fluorescently labeled with 5(6)-carboxyfluorescein was prepared in n-octylglucoside as previously described for immunoblot analyses. Briefly, UMSCC2 cells were seeded on 22 mm × 22 mm chambered coverslips pretreated with poly-l-lysine at 300,000 cells/well to be ~60% confluent after 16 h. The cells were then treated with 10 μM peptide in complete media at 37 °C for 1 h. The cells were washed with PBS before being imaged by spinning disc confocal microscopy.

## Acknowledgements

We thank Prof. Aurelia Honerkamp-Smith (Department of Physics, Lehigh University) for technical assistance and use of the spinning disk microscope. This work was supported by the National Cancer Institute [grant number R21CA195158] to D.T. and M.J.L.; and start-up funds from Lehigh University to D.T.

## Author Contributions

D.T. and M.J.L. designed research; E.B., E.S., D.P., and E.K.D. performed research; B.W.B. designed the peptide binder library; E.B., E.S., M.J.L, and D.T. analyzed the data; E.B., E.S., M.J.L, and D.T. wrote the paper.

## Abbreviations

BSA: Bovine serum albumin
CD: Circular Dichroism
EGFR: Epidermal Growth Factor Receptor
EV: Empty Vector
GFP: Green Fluorescence Protein
MALDI-TOF: Matrix-assisted laser desorption/ionization time of flight
PLA: Proximity Ligation Assay
POPC: 1-palmitoyl-2-oleoyl-sn-glycero-3-phosphocholine
TM: Transmembrane
TBS-T’: Tris-buffered saline Tween 20
RPTP: Receptor Protein Tyrosine Phosphatase
RT: Room Temperature
RTK: Receptor Tyrosine Kinase
WT: Wild-type

## References

1. Barr, A. J., Ugochukwu, E., Lee, W. H., King, O. N. F., Filippakopoulos, P., Alfano, I., Savitsky, P., Burgess-Brown, N. A., Müller, S., and Knapp, S. (2009) Large-Scale Structural Analysis of the Classical Human Protein Tyrosine Phosphatome. Cell. 136, 352–363

2. Tonks, N. K. (2006) Protein tyrosine phosphatases: from genes, to function, to disease. Nat Rev Mol Cell Biol. 7, 833–846

3. Tonks, N. K. (2013) Protein tyrosine phosphatases--from housekeeping enzymes to master regulators of signal transduction. FEBS J. 280, 346–378

4. Ruivenkamp, C., Hermsen, M., Postma, C., Klous, A., Baak, J., Meijer, G., and Demant, P. (2003) LOH of PTPRJ occurs early in colorectal cancer and is associated with chromosomal loss of 18q12-21. Oncogene. 22, 3472–3474

5. Ruivenkamp, C. A. L., van Wezel, T., Zanon, C., Stassen, A. P. M., Vlcek, C., Csikós, T., Klous, M., Tripodis, N., Perrakis, A., Boerrigter, L., Groot, P. C., Lindeman, J., Mooi, W. J., Meijjer, G. A., Scholten, G., Dauwerse, H., Paces, V., van Zandwijk, N., van Ommen, G. J. B., and Demant, P. (2002) Ptprj is a candidate for the mouse colon-cancer susceptibility locus Scc1 and is frequently deleted in human cancers. Nat Genet. 31, 295–300

6. Nunes-Xavier, C. E., Martín-Pérez, J., Elson, A., and Pulido, R. (2013) Protein tyrosine phosphatases as novel targets in breast cancer therapy. Biochim Biophys Acta. 1836, 211–226

7. Iuliano, R., Le Pera, I., Cristofaro, C., Baudi, F., Arturi, F., Pallante, P., Martelli, M. L., Trapasso, F., Chiariotti, L., and Fusco, A. (2004) The tyrosine phosphatase PTPRJ/DEP-1 genotype affects thyroid carcinogenesis. Oncogene. 23, 8432–8438

8. Keane, M. M., Lowrey, G. A., Ettenberg, S. A., Dayton, M. A., and Lipkowitz, S. (1996) The protein tyrosine phosphatase DEP-1 is induced during differentiation and inhibits growth of breast cancer cells. Cancer Res. 56, 4236–4243

9. Ostman, A., Hellberg, C., and Böhmer, F.-D. (2016) Protein Tyrosine Phosphatases in Cancer (Neel, B. G., and Tonks, N. K. eds), Springer-Verlag New York, 6, 307–320

10. Tarcic, G., Boguslavsky, S. K., Wakim, J., Kiuchi, T., Liu, A., Reinitz, F., Nathanson, D., Takahashi, T., Mischel, P. S., Ng, T., and Yarden, Y. (2009) An unbiased screen identifies DEP-1 tumor suppressor as a phosphatase controlling EGFR endocytosis. Curr Biol. 19, 1788–1798

11. Avraham, R., and Yarden, Y. (2011) Feedback regulation of EGFR signalling: decision making by early and delayed loops. Nat Rev Mol Cell Biol. 12, 104–117

12. Citri, A., and Yarden, Y. (2006) EGF-ERBB signalling: towards the systems level. Nat Rev Mol Cell Biol. 7, 505–516

13. Zhang, L., Martelli, M. L., Battaglia, C., Trapasso, F., Tramontano, D., Viglietto, G., Porcellini, A., Santoro, M., and Fusco, A. (1997) Thyroid cell transformation inhibits the expression of a novel rat protein tyrosine phosphatase. Exp. Cell Res. 235, 62–70

14. Stanford, S. M., and Bottini, N. (2017) Targeting Tyrosine Phosphatases: Time to End the Stigma. Trends Pharmacol Sci. 38, 524–540

15. Jiang, G., Hertog, den, J., Su, J., Noel, J., Sap, J., and Hunter, T. (1999) Dimerization inhibits the activity of receptor-like protein-tyrosine phosphatase-alpha. Nature. 401, 606–610

16. Bilwes, A. M., Hertog, den, J., Hunter, T., and Noel, J. P. (1996) Structural basis for inhibition of receptor protein-tyrosine phosphatase-alpha by dimerization. Nature. 382, 555–559

17. Tertoolen, L. G., Blanchetot, C., Jiang, G., Overvoorde, J., Gadella, T. W., Hunter, T., and Hertog, den, J. (2001) Dimerization of receptor protein-tyrosine phosphatase alpha in living cells. BMC Cell Biol. 2, 8

18. Chin, C.-N., Sachs, J. N., and Engelman, D. M. (2005) Transmembrane homodimerization of receptor-like protein tyrosine phosphatases. FEBS Lett. 579, 3855–3858

19. Takeda, A., Matsuda, A., Paul, R. M. J., and Yaseen, N. R. (2004) CD45-associated protein inhibits CD45 dimerization and up-regulates its protein tyrosine phosphatase activity. Blood. 103, 3440–3447

20. Cahir McFarland, E. D., Pingel, J., and Thomas, M. L. (1997) Definition of amino acids sufficient for plasma membrane association of CD45 and CD45-associated protein. 36, 7169–7175

21. Su, P.-C., and Berger, B. W. (2012) Identifying key juxtamembrane interactions in cell membranes using AraC-based transcriptional reporter assay (AraTM). J Biol Chem. 287, 31515–31526

22. Su, P.-C., and Berger, B. W. (2013) A novel assay for assessing juxtamembrane and transmembrane domain interactions important for receptor heterodimerization. 425, 4652–4658

23. MacKenzie, K. R., and Engelman, D. M. (1998) Structure-based prediction of the stability of transmembrane helix-helix interactions: the sequence dependence of glycophorin A dimerization. Proc Natl Acad Sci USA. 95, 3583–3590

24. Kim, S., Jeon, T.-J., Oberai, A., Yang, D., Schmidt, J. J., and Bowie, J. U. (2005) Transmembrane glycine zippers: physiological and pathological roles in membrane proteins. Proc Natl Acad Sci USA. 102, 14278–14283

25. Russ, W. P., and Engelman, D. M. (2000) The GxxxG motif: a framework for transmembrane helix-helix association. 296, 911–919

26. Senes, A., Engel, D. E., and DeGrado, W. F. (2004) Folding of helical membrane proteins: the role of polar, GxxxG-like and proline motifs. Curr Opin Struct Biol. 14, 465–479

27. Mueller, B. K., Subramaniam, S., and Senes, A. (2014) A frequent, GxxxG-mediated, transmembrane association motif is optimized for the formation of interhelical Cα-H hydrogen bonds. Proc. Natl. Acad. Sci. U.S.A. 111, E888–95

28. Langosch, D. (2015) Role of GxxxG Motifs in Transmembrane Domain Interactions. Biochemistry. 54, 5125–5135

29. Böhmer, S.-A., Weibrecht, I., Söderberg, O., and Böhmer, F.-D. (2013) Association of the Protein-Tyrosine Phosphatase DEP-1 with Its Substrate FLT3 Visualized by In Situ Proximity Ligation Assay. PLoS ONE. 8, e62871

30. Mellberg, S., Dimberg, A., Bahram, F., Hayashi, M., Rennel, E., Ameur, A., Westholm, J. O., Larsson, E., Lindahl, P., Cross, M. J., and Claesson-Welsh, L. (2009) Transcriptional profiling reveals a critical role for tyrosine phosphatase VE-PTP in regulation of VEGFR2 activity and endothelial cell morphogenesis. FASEB J. 23, 1490–1502

31. Hayashi, M., Majumdar, A., Li, X., Adler, J., Sun, Z., Vertuani, S., Hellberg, C., Mellberg, S., Koch, S., Dimberg, A., Koh, G. Y., Dejana, E., Belting, H.-G., Affolter, M., Thurston, G., Holmgren, L., Vestweber, D., and Claesson-Welsh, L. (2013) VE-PTP regulates VEGFR2 activity in stalk cells to establish endothelial cell polarity and lumen formation. Nat Commun. 4, 1672

32. Heim, E. N., Marston, J. L., Federman, R. S., Edwards, A. P. B., Karabadzhak, A. G., Petti, L. M., Engelman, D. M., and DiMaio, D. (2015) Biologically active LIL proteins built with minimal chemical diversity. Proc. Natl. Acad. Sci. U.S.A. 112, E4717–25

33. Freeman-Cook, L. L., Dixon, A. M., Frank, J. B., Xia, Y., Ely, L., Gerstein, M., Engelman, D. M., and DiMaio, D. (2004) Selection and characterization of small random transmembrane proteins that bind and activate the platelet-derived growth factor beta receptor. Journal of Molecular Biology. 338, 907–920

34. Goh, E. T. H., Lin, Z., Ahn, B. Y., Lopes-Rodrigues, V., Dang, N. H., Salim, S., Berger, B., Dymock, B., Senger, D. L., and Ibáñez, C. F. (2018) A Small Molecule Targeting the Transmembrane Domain of Death Receptor p75NTR Induces Melanoma Cell Death and Reduces Tumor Growth. Cell Chem Biol. 25, 1485–1494.e5

35. Thévenin, D., and Lazarova, T. (2012) Identifying and measuring transmembrane helix-helix interactions by FRET. Methods Mol Biol. 914, 87–106

36. Arpel, A., Sawma, P., Spenlé, C., Fritz, J., Meyer, L., Garnier, N., Velázquez-Quesada, I., Hussenet, T., Aci-Sèche, S., Baumlin, N., Genest, M., Brasse, D., Hubert, P., Crémel, G., Orend, G., Laquerrière, P., and Bagnard, D. (2014) Transmembrane domain targeting peptide antagonizing ErbB2/Neu inhibits breast tumor growth and metastasis. Cell Rep. 8, 1714–1721

37. Nasarre, C., Roth, M., Jacob, L., Roth, L., Koncina, E., Thien, A., Labourdette, G., Poulet, P., Hubert, P., Crémel, G., Roussel, G., Aunis, D., and Bagnard, D. (2010) Peptide-based interference of the transmembrane domain of neuropilin-1 inhibits glioma growth in vivo. Oncogene. 29, 2381–2392

38. Hubert, P., Sawma, P., Duneau, J.-P., Khao, J., Hénin, J., Bagnard, D., and Sturgis, J. (2010) Single-spanning transmembrane domains in cell growth and cell-cell interactions: More than meets the eye? Cell Adh Migr. 4, 313–324

39. Bennasroune, A., Fickova, M., Gardin, A., Dirrig-Grosch, S., Aunis, D., Crémel, G., and Hubert, P. (2004) Transmembrane peptides as inhibitors of ErbB receptor signaling. Mol Biol Cell. 15, 3464–3474

40. Iuliano, R., Raso, C., Quintiero, A., Pera, I. L., Pichiorri, F., Palumbo, T., Palmieri, D., Pattarozzi, A., Florio, T., Viglietto, G., Trapasso, F., Croce, C. M., and Fusco, A. (2009) The Eighth Fibronectin Type III Domain of Protein Tyrosine Phosphatase Receptor J Influences the Formation of Protein Complexes and Cell Localization. Journal of Biochemistry. 145, 377–385

41. Monast, C. S., Furcht, C. M., and Lazzara, M. J. (2012) Computational analysis of the regulation of EGFR by protein tyrosine phosphatases. Biophys J. 102, 2012–2021

42. Whiteford, J. R., Xian, X., Chaussade, C., Vanhaesebroeck, B., Nourshargh, S., and Couchman, J. R. (2011) Syndecan-2 is a novel ligand for the protein tyrosine phosphatase receptor CD148. Mol Biol Cell. 22, 3609–3624

43. Takahashi, K., Mernaugh, R. L., Friedman, D. B., Weller, R., Tsuboi, N., Yamashita, H., Quaranta, V., and Takahashi, T. (2012) Thrombospondin-1 acts as a ligand for CD148 tyrosine phosphatase. Proc. Natl. Acad. Sci. U.S.A. 109, 1985–1990

44. Grazia Lampugnani, M., Zanetti, A., Corada, M., Takahashi, T., Balconi, G., Breviario, F., Orsenigo, F., Cattelino, A., Kemler, R., Daniel, T. O., and Dejana, E. (2003) Contact inhibition of VEGF-induced proliferation requires vascular endothelial cadherin, beta-catenin, and the phosphatase DEP-1/CD148. The Journal of Cell Biology. 161, 793–804

45. Chabot, C., Spring, K., Gratton, J.-P., Elchebly, M., and Royal, I. (2009) New role for the protein tyrosine phosphatase DEP-1 in Akt activation and endothelial cell survival. Mol Cell Biol. 29, 241–253

46. Takahashi, T., Takahashi, K., Mernaugh, R. L., Tsuboi, N., Liu, H., and Daniel, T. O. (2006) A monoclonal antibody against CD148, a receptor-like tyrosine phosphatase, inhibits endothelial-cell growth and angiogenesis. Blood. 108, 1234–1242

47. Palka, H. L., Park, M., and Tonks, N. K. (2003) Hepatocyte growth factor receptor tyrosine kinase met is a substrate of the receptor protein-tyrosine phosphatase DEP-1. J Biol Chem. 278, 5728–5735

48. Xu, Y., Xia, W., Baker, D., Zhou, J., Cha, H. C., Voorhees, J. J., and Fisher, G. J. (2011) Receptor-type protein tyrosine phosphatase beta (RPTP-beta) directly dephosphorylates and regulates hepatocyte growth factor receptor (HGFR/Met) function. J Biol Chem. 286, 15980–15988

49. Sacco, F., Tinti, M., Palma, A., Ferrari, E., Nardozza, A. P., Hooft van Huijsduijnen, R., Takahashi, T., Castagnoli, L., and Cesareni, G. (2009) Tumor suppressor density-enhanced phosphatase-1 (DEP-1) inhibits the RAS pathway by direct dephosphorylation of ERK1/2 kinases. J Biol Chem. 284, 22048–22058

