## Supplementary Figures for "Disrupting the Transmembrane Domain Oligomerization of Protein Tyrosine Phosphatase Receptor J Promotes its Activity in Cancer Cells"

- **Fig. S1.** Maltose complementation test.
- **Fig. S2.** Expression level of PTPR and EGFR.
- **Fig. S3.** Peptides purity check and mass spectrometry
- **Fig. S4.** Interaction of $RJ_{binder}$ and $RJ_{control}$ with membrane mimics by circular dichroism and fluorescence spectroscopies
- **Fig. S5.** $RJ_{binder}$ partitions into cell membranes by fluorescence microscopy
- **Fig. S6.** $RJ_{binder}$ has no effect on migration in cells expressing minimal PTPRJ (EV) or the homodimer-disrupting mutant (G983L).
- **Fig. S7.** Sensitivity of UMSCC2 cells expressing WT PTPRJT towards gefitinib.

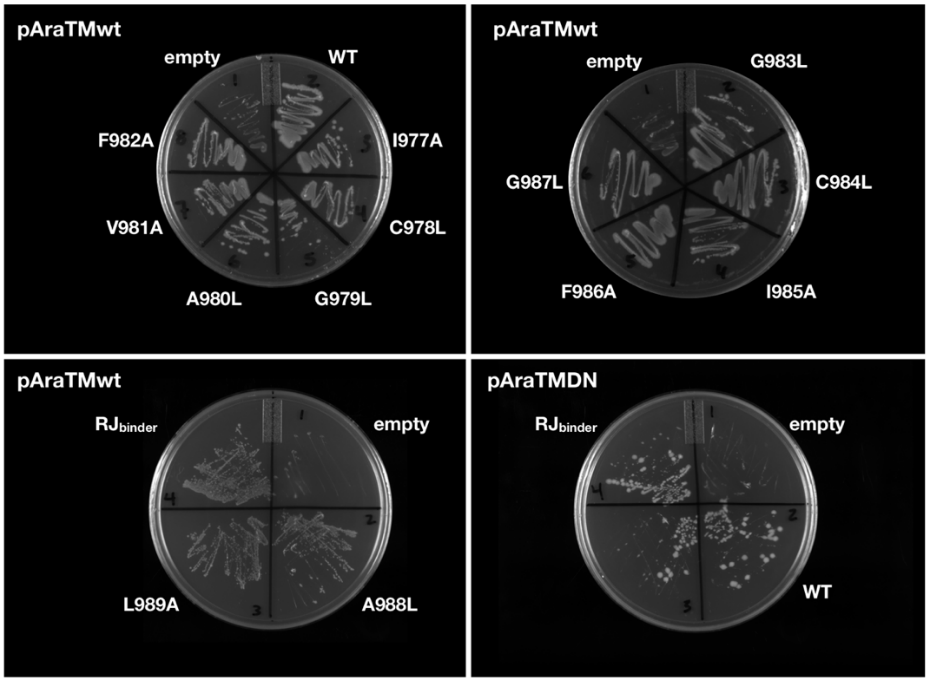


**Fig. S1. Maltose complementation test.** The indicated pAraTMwt or pAraTMDN constructs were transformed into the MBP-deficient *E. coli* strain MM39. Saturated culture from each chimera was streaked onto M9 minimal media plates containing maltose. No growth was observed in the empty vectors, in which no AraC or AraC* fusion is expressed. Recovery of growth was observed for all other fusion constructs, indicating proper orientation in the cell membrane.

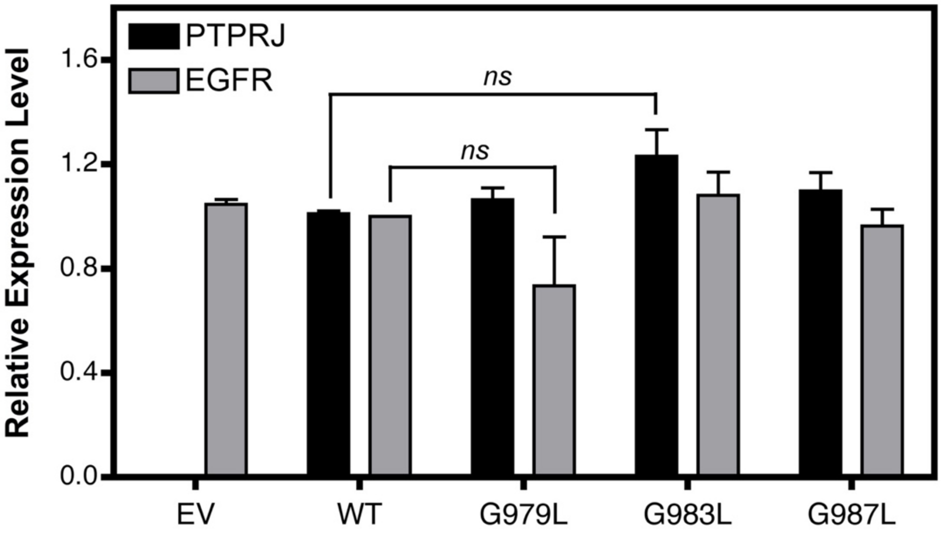


**Fig. S2.** Relative expression levels of total PTPRJ and EGFR in UMSCC2 cells normalized to WT. Results are shown as mean ± SEM ($n$ = 3). Statistical significance was assessed using unpaired $t$ test (at 95% confidence intervals).

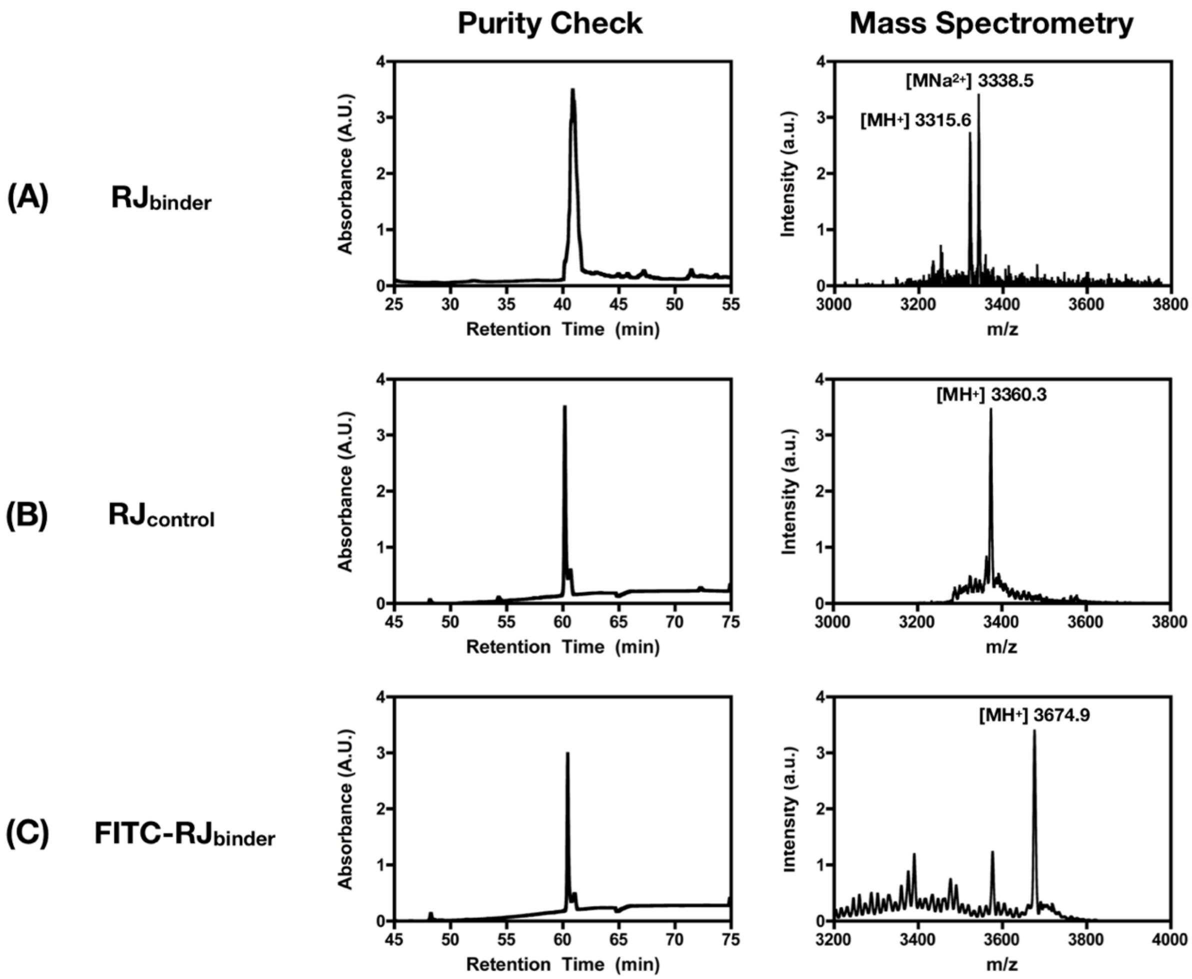


**Fig. S3.** Purity check by RP-HPLC and MALDI-TOF MS spectra of synthesized peptides. **(A)** $RJ_{binder}$: calculated ($MH^+$) = 3316.1 g/mol, found ($MH^+$) 3315.6 g/mol. **(B)** $RJ_{control}$: calculated ($MH^+$) 3358.2, found ($MH^+$) = 3360.3 g/mol. **(C)** FITC-$RJ_{binder}$: calculated ($MH^+$) = 3674.5 g/mol, found ($MH^+$) = 3674.9 g/mol.

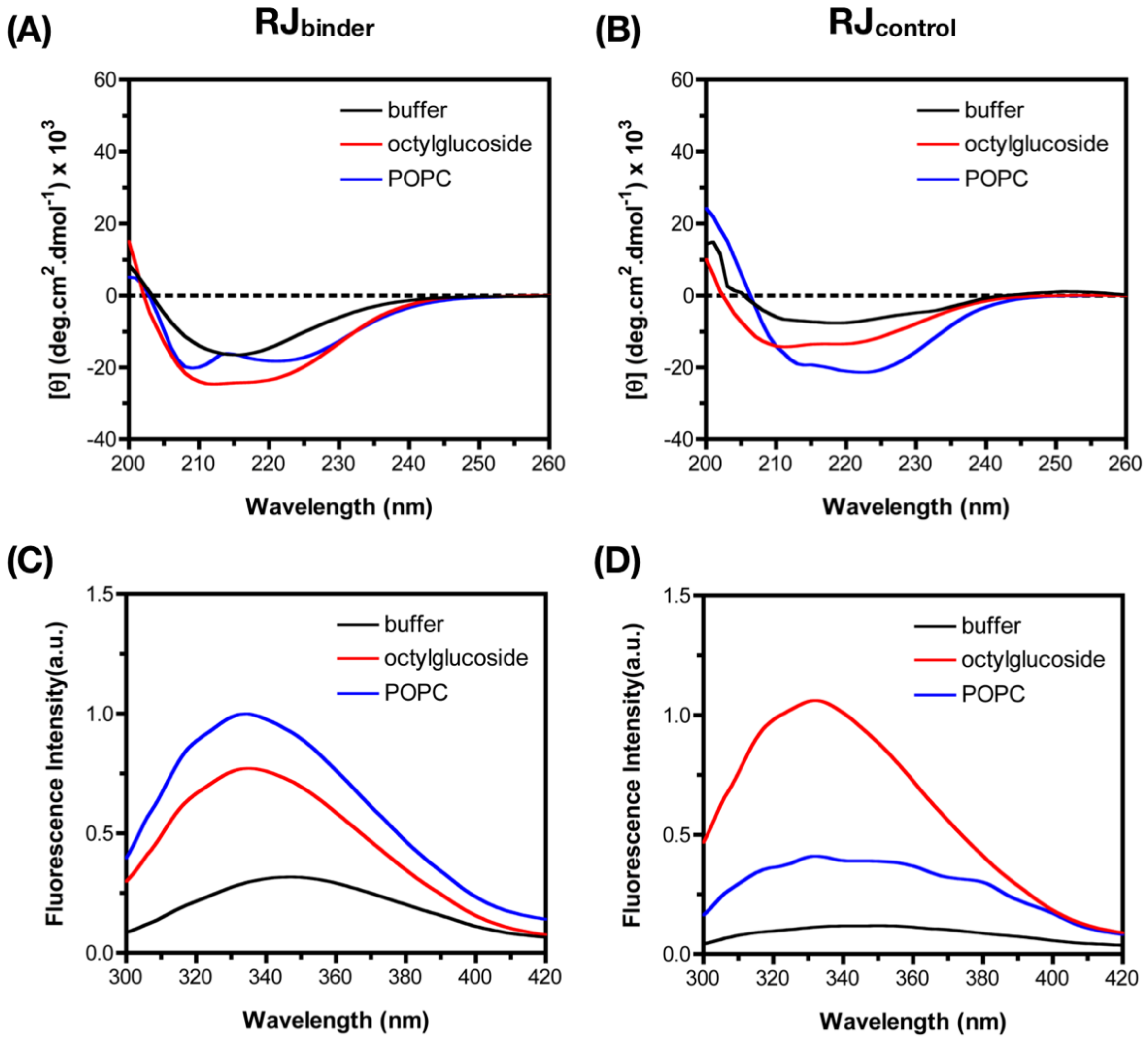


**Fig. S4. Interaction of RJ$_{binder}$ and RJ$_{control}$ with membrane mimics**. Circular dichroism **(A,B)** and tryptophan fluorescence emission **(C,D)** of 10 μM RJ$_{binder}$ and RJ$_{control}$ in buffer (black), 30 mM n-octylglucoside micelles (red) or large unilamellar POPC lipid vesicles at a 1:300 peptide/lipid ratio (blue).

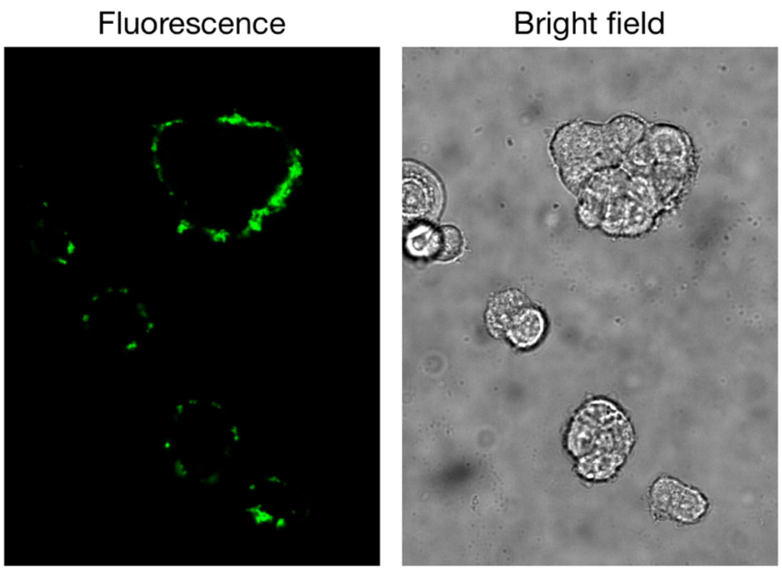


**Fig. S5. RJ$_{binder}$ partitions into cell membranes.** UMSCC2 cells were treated with 10 μM fluorescently labeled RJ$_{binder}$ reconstituted in n-octyliglucoside micelles for 1 h. Representative fluorescence and bright field images taken from spinning disk confocal microscopy are shown.

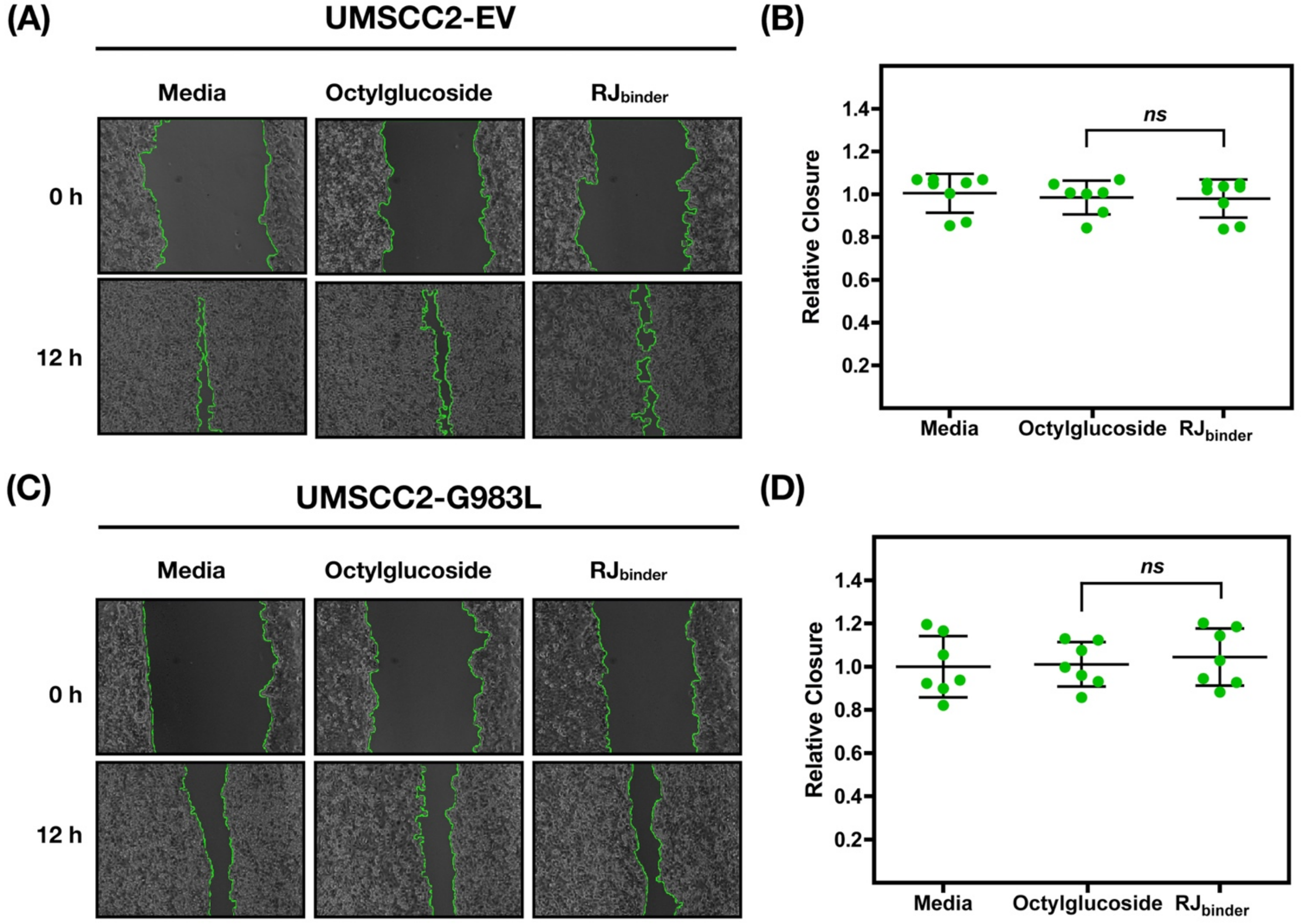


**Fig. S6. RJ$_{binder}$ has no effect on migration in cells expressing minimal PTPRJ (EV) or the homodimer-disrupting mutant (G983L).** Representative phase contrast images with tracings to identify open scratch areas **(A,C)** and quantification **(B,D)** of the effect of RJ$_{binder}$ on cells expressing either minimal PTPRJ **(A,B)** or G983L PTPRJ **(C,D)**. Serum-starved UMSCC2 cells were treated with 10 µM RJ$_{binder}$ reconstituted in n-octylglucoside micelles for 1 h, scratched (0 h), and incubated media containing EGF (50 ng/mL) for 12 h. Relative closure was quantified by calculating the percent change in area between 0 and 12 h using ImageJ, and then normalized to media containing EGF. **(B,D)** Results are shown as mean ± SEM ($n$ = 6-9). Statistical significance was assessed using unpaired $t$ test (at 95% confidence intervals).

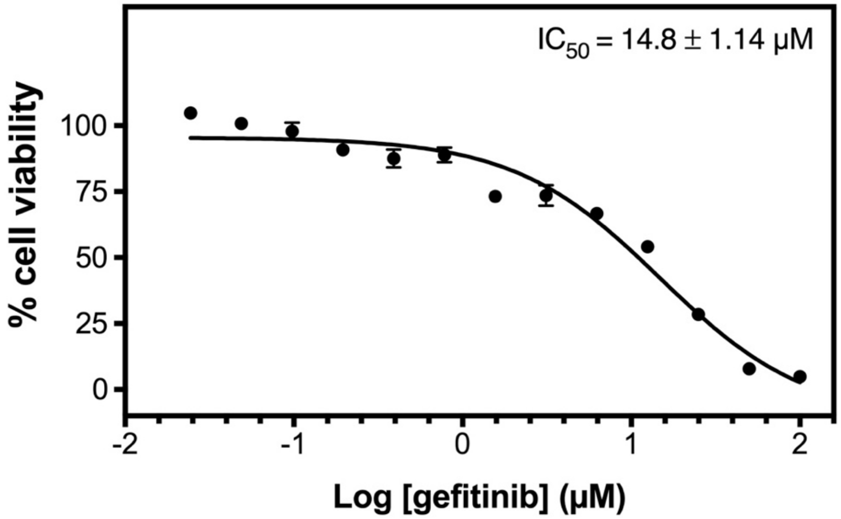


**Fig. S7. Sensitivity of UMSCC2 cells expressing WT PTPRJT towards gefitinib.** 5,000 cells/well were treated with increasing concentrations of gefitinib in 1% DMSO for 72 hours. Cell viability was determined using the colorimetric MTT assay. Briefly, 10 µL of a 5 mg/mL MTT stock solution was added to the treated cells and incubated for 2 h at 37 °C. The resulting formazan crystals were solubilized in 200 µL DMSO and the absorbance measured at 580 nm using an Infinite 200 PRO microplate reader (Teca). Results are shown as mean ± SEM (*n* = 6). Data were fitted with a sigmoidal dose-response (Prism for Mac, GraphPad, Inc.).
